# Hierarchical behavior control by a single class of interneurons

**DOI:** 10.1101/2023.03.13.532397

**Authors:** Jing Huo, Tianqi Xu, Qi Liu, Mahiber Polat, Sandeep Kumar, Xiaoqian Zhang, Andrew M. Leifer, Quan Wen

## Abstract

Animal behavior is organized into nested temporal patterns that span multiple timescales. This behavior hierarchy is believed to arise from a hierarchical neural architecture: neurons near the top of the hierarchy are involved in planning, selecting, initiating, and maintaining motor programs, whereas those near the bottom of the hierarchy act in concert to produce fine spatiotemporal motor activity. In *Caenorhabditis elegans*, behavior on a long timescale emerges from ordered and flexible transitions between different behavioral states, such as forward, reversal, and turn. On a short timescale, different parts of the animal body coordinate fast rhythmic bending sequences to produce directional movements. Here, we show that SAA, a class of interneurons that enable cross-communication between dorsal and ventral head motor neurons, play a dual role in shaping behavioral dynamics on different timescales. On a short timescale, SAA regulate and stabilize rhythmic bending activity during forward movements. On a long timescale, the same neurons suppress spontaneous reversals and facilitate reversal termination by inhibiting RIM, an integrating neuron that helps maintain a behavioral state. These results suggest that feedback from a lower-level cell assembly to a higher-level command center is essential for bridging behavioral dynamics at different levels.

**Significance Statement:** In this study, we reveal the dual role of SAA interneurons in *C. elegans*, demonstrating their influence over diverse behavior timescales. These neurons not only stabilize short-term rhythmic activities during forward movements, but also modulate long-term behavioral transitions between motor states. This indicates the essential role of feedback from low-level neural assemblies to command centers in a hierarchical neural architecture, emphasizing its significance in orchestrating behavior across scales. Our study offers critical insights into the intricate neural interactions behind organized and adaptive behavior.

---

In his book *The Study of Instinct*, Tinbergen proposed that animal behavior is organized into a hierarchical structure of muscle activity patterns (1). Nest-building in stickleback, for example, is composed of a series of behaviors - like digging, testing materials, boring, and gluing — each of which can be subdivided into finer actions (on pages 133-134, (1)). The nested structures across multiple timescales is the basis of behavioral hierarchy, where a behavioral module spanning a longer timescale stays near the top of the hierarchy. This theory has been elaborated and tested in classic ethological studies (2–4) and more recently by modern machine learning approaches (5–10). In particular, an impartial method for classifying behaviors and quantifying their relationships reveals a remarkable tree-like structure among all observable behavioral motifs in *Drosophila* (7).

The behavioral hierarchy is believed to originate from a hierarchical neural architecture of movement control (11–14): the neural code that represents a specific behavioral state is sparse and organized centrally, while the neural representation that contributes to muscle synergy is distributed towards the periphery with increasingly dense and fast dynamics (15). Studies from several animal models have appeared to support this view. During the singing of zebra finches, a projection neuron in a premotor nucleus HVC generates a sparse and short burst of spikes reliably at one precise moment in the entire song sequence (16, 17), while neurons in the downstream nucleus RA, which project to motor neurons to control the syrinx, exhibit dense and variable firing (18). The foraging state in larval zebrafish, during which the animal suppresses swimming and promotes hunting, is represented by persistent activity in a sparse neural population in dorsal raphe (19). Hunting behavior, which is composed of rapid eye convergence and body J-turn, is represented by fast brain-wide activity across many midbrain and hindbrain areas (20). The attacking state in the mouse is represented by slowly varying population neural activity in a VMHv nucleus in the hypothalamus, while sequential actions involving faster dynamics are encoded by different groups of neurons in a downstream nucleus MPOA (21).

The neural mechanisms underlying the nested temporal patterns in naturalistic behavior, however, are poorly understood. The nematode *Caenorhabditis elegans* offers an opportunity to develop a deep understanding of the hierarchy problem, since the neural basis of worm behavior on different timescales has been studied in great detail (13, 22–31). Let us use foraging behavior as an example.

- At the top level, the *C. elegans* exploratory behavior exhibits two different strategies: local search shortly after the animals are removed from food and global dispersal after prolonged food deprivation (24). In the presence of food, the *C. elegans* foraging behavior also exhibits two similar behavior states, namely dwelling and roaming (32). Two groups of neurons, NSM/HSN and PVP/AVB, which release serotonin and the neuropeptide PDF, respectively, were shown to play opposing roles in modulating the dwelling and roaming state.
- At the middle level, *C. elegans* locomotion consists of forward movement, reversal, and turn. Local search is associated with a higher frequency of reversals and turns, whereas global dispersal promotes forward movements and inhibits reversals. Recent Ca^2+^ imaging of the neuronal population in immobilized and freely-behaving animals revealed that persistent activities in different groups of interneurons represent distinct motor states (29, 33, 34). For example, AVB/AIY/RIB exhibit elevated calcium activity during forward movements, while AVA/RIM/AIB exhibit elevated activity during reversal. The ordered and flexible sequential transitions between behaviors are controlled by a combination of excitatory and inhibitory interactions between cell assemblies and a winner-take-all strategy for action selection (35–38).
- At the bottom level, directional movements require fast rhythmic bending waves that propagate throughout the worm body. Forward movement in *C. elegans* is driven by B-type motor neurons and head motor neurons while reversal is driven by A-type motor neurons. Descending input from AVB, for example, is critical for triggering rhythmic activity in midbody B-type motor neurons (30, 39). During reversal, AVA promotes rhythmic activity in A-type motor neurons (26, 37), while AIB/RIM inhibit SMD motor neurons to suppress head movements (13, 40).

The mounting experimental evidence suggests a framework (fig. 1) for organizing behavior across timescales: neural activity at each level along the hierarchy can have its intrinsic timescale, determined by the biophysical properties of neurons, the interactions between neurons within the same group, and the influence of neuromodulators. Feedforward inputs from top to bottom layers are primarily involved in selecting and gating diverse temporal patterns. This raises the intriguing question of whether this structure is purely feedforward. This view is challenged by two earlier studies. In particular, Kaplan et al. (13) showed that continuous inhibition of SMD head motor neurons through exogenous expression of the histamine-chloride channel not only curbed head bending motor activity during forward movements but also prolonged reversal duration. Optogenetic inhibition of a small portion of B-type motor neurons in the midbody of the animal could render the entire animal immobile under high-light intensity illumination or decelerate movement under low-light intensity illumination (30). These findings suggest the existence of retrograde signals to reconfigure the dynamics of the motor circuit on various timescales.

**Fig. 1.**
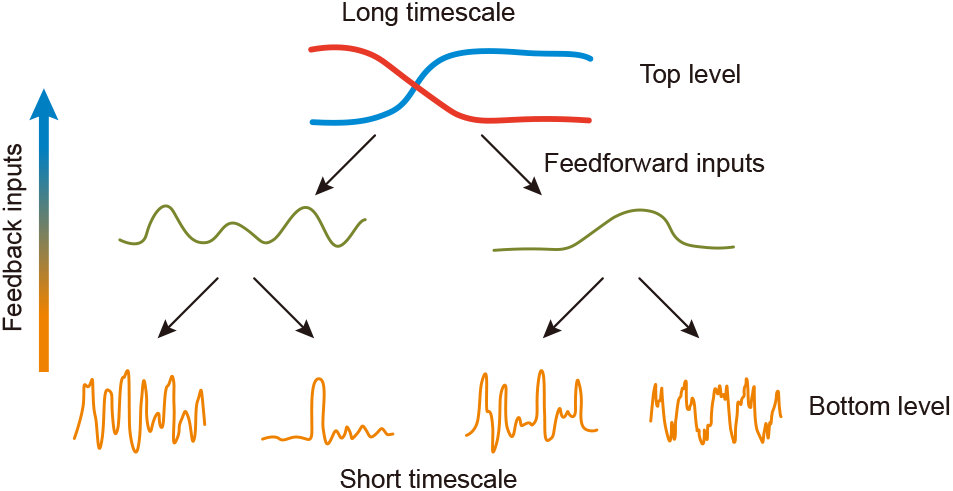
A conceptual scheme for organizing behaviors across different timescales. Top level: two temporal sequences illustrate the neural activity of two separate circuits, each resulting in unique behavioral strategies. The prevailing strategy can affect neural activity at subordinate levels across extended timescales. Bottom level: the neural activity or behavioral patterns of motor systems that implement the prevailing strategy. Circuits located at the upper level send feedforward instructions to the lower level, whereas lower level circuits have the capability to send feedback signals to the upper level.

Inspired by earlier work (7, 13, 41), here we show that a class of interneurons SAA within the *C. elegans* head motor circuit play a dual role in shaping the timescales of low-level and high-level behavior dynamics. SAA make numerous connections with dorsal and ventral head motor neurons SMB/SMD/RMD, through either by gap junctions or chemical synapses. We demonstrate that, on a short timescale, SAA regulate head-bending kinematics and coordinate undulatory wave propagation during forward movements. Remarkably, on the long timescale, we find that feedback inhibition from SAA to RIM, an integrating neuron in the motor state control center, facilitates reversal termination and impacts stochastic transitions between motor states. Feedback from a lower-level to a higher-level circuitry (fig. 1) *complements voluntary control* that uses a strict top-down strategy; and we argue that the presence of loops in a *pyramid-like* architecture (2) provides a more efficient and robust way to control behavior.

## Results

### SAA regulate and stabilize fast kinematics in forward locomotion

The *C. elegans* connectome (41, 42) indicates that SAA play a special role in the head motor circuit. SAA represents four interneurons (SAADL/R, SAAVL/R) that make reciprocal connections, either by electrical synapses or chemical synapses, with three classes of head motor neurons, SMB/SMD/RMD. For example, each of the four SMB motor neurons sends synapses to SAA, and SAA, in turn, makes gap junctions with the SMB neurons that innervate muscles on the opposite side (fig. 2A). This circuit motif allows for contralateral communication between SMBV and SMBD, which are not directly connected to each other.

**Fig. 2.**
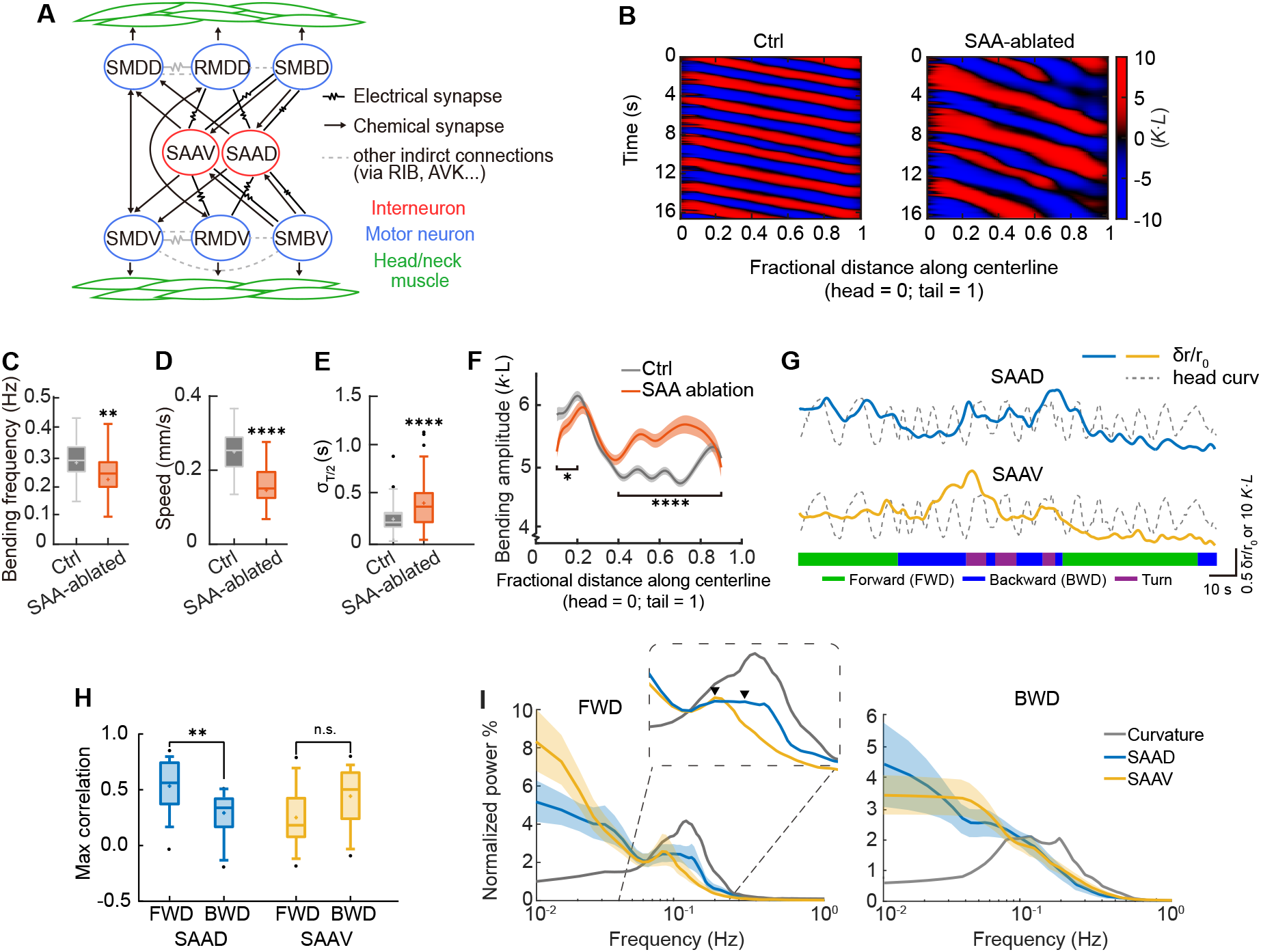
SAA neurons modulate fast kinematics in forward locomotion. **A**. Diagrams of circuit motif featuring SAAV and SAAD together with head motor neurons. **B**. Representative curvature kymographs for control (mock-ablated) and SAA-ablated animals during forward locomotion. Body curvature is expressed as a dimensionless unit *κ* · *L*, normalized by the worm’s body length *L*. **C-E**. Forward movement velocity, undulation frequency and standard deviation of the half-cycle duration (see text). **P = 0.001, ****P *<* 0.0001, two-sample t-test with Welch’s correction. Box plots show the quartiles of the dataset, with whiskers extending from minimum to maximum, black dots are outliers and cross signs are mean. Ctrl: n = 113 trials (episodes), 8 animals; SAA-ablated: n = 85 trials, 8 animals. Both the control group and the actual SAA-ablated group consist of transgenic animals (P*lad-2*::Cre; P*lim-4*::loxP::PH-miniSOG). **F**. Whole body bending amplitude for both control and SAA-ablated animals. The line indicates the average; the shaded region represents the SEM across trials. *P = 0.016, fractional distance *∈* [0.1, 0.2]; ****P *<* 0.0001, fractional distance *∈* [0.4, 0.9], Mann–Whitney U test was performed on the spatially averaged bending amplitude across the body of the worm (Methods). Ctrl: n = 113, 8 animals; SAA-ablated: n = 85, 8 animals. **G**. The representative trace displays the activity of SAAD/V together with the curvature of the worm’s head during locomotion. A blue trace illustrates the change in the ratio between GECI wNEMOs (43) and wCherry. A dark dashed trace indicates the dynamics of head curvature ([0.1,0.2] fractional distance along the worm, see **A**), with positive values showing dorsal bending of the head. The colored ribbons below depict the behavior states throughout 140-second traces. The worm moved on a 2% agarose pad covered with a glass slide. **H**. Maximum correlation between the neuronal activity and head curvature during forward or backward locomotion. We aligned the neuronal signals with a time-shifted (within *±* 2 s) head curvatures to find the highest cross-correlation. **P = 0.005, for SAAD result, n.s. p=0.073, using a two-sample t-test with Welch’s adjustment. The box plots illustrate the quartiles of the data, with strike bars indicating the range from minimum to maximum, black dots marking outliers, and cross signs denoting average. n=16 for forward movement and n=12 for backward movement, across 8 animals. **I**. Average power spectrum density for SAAD and SAAV during forward or backward locomotion. The peak of the PSD curves are indicated by triangles. The shaded regions show the standard deviations. For forward movement, n = 16 and for backward movement, n = 12, from a total of 8 worms.

To determine the functional contribution of SAA to head motor activity and *C. elegans* locomotion, we generated transgenic animals (P*lad-2*::Cre; P*lim-4*::loxP::PH-miniSOG) that specifically enable optogenetic ablation of SAA neurons (Methods). The kinematics of the worm bending activity during forward movements on an agarose pad can be visualized by a curvature kymograph: in a control animal, each body segment alternated between positive (red) and negative (blue) curvature, and the curvature bands propagated regularly from head to tail (head = 0; tail = 1) (fig. 2B). In an SAA-ablated animal, the curvature bands appeared wider; in addition, we found that the undulation frequency (fig. 2C, fig. S2A) as well as the speed of locomotion (fig. 2D, fig. S2B) were significantly reduced. In line with the findings from the ablation experiments, rapid optogenetic inhibition of SAA significantly decreased the speed and frequency of forward movement (fig. S1A,D), but the effects were weaker than SAA ablation. Optogenetic stimulation of SAA, however, did not cause significant alterations in forward movement kinematics.

Interestingly, the deceleration of movements in SAA-ablated animals was associated with increased rhythmic irregularity. When crawling, wild-type animals generate a trajectory similar to that of a sinusoid. We determined the body curvature (fig. 2B) and computed the temporal difference between consecutive peaks and troughs, a measure we refer to as the semiperiod. This semiperiod showed substantial variation over time in SAA-ablated or inhibited animals, as indicated by a higher standard deviation (SD) (σ _*T/*2_ in fig. 2E, fig. S1B), leading to a broader distribution. A similar change in the distribution of the semiperiod was observed in swimming animals (fig. S2D). The head bending amplitude was reduced, whereas the bending amplitude in the mid-body significantly increased (fig. 2F).

The spatiotemporal correlation of the bending activity (fig. 2B) also suggests that the dynamic motion of an entire animal can be described by a small number of collective variables using Principal Component Analysis (PCA) (25), and the time evolution of worm behavior can be recapitulated by a trajectory in a low-dimensional phase space ((44) and Methods). A typical phase trajectory of a control animal appeared circular (fig. 3A), indicating regular periodic motion; the phase trajectory of an SAA-ablated animal, however, was broadly extended (fig. 3B). By examining the density of the phase trajectories across trials and animals (fig. 3C, D), we found that the motions of control animals were restricted to a smaller region similar to a torus, while those of SAA-ablated animals were more broadly dispersed in the phase space (fig. 3E). A similar change in the shape of the trajectories was also observed in swimming animals (fig. S3). We further evaluated control animals moving at low speeds comparable to SAA-ablated animals (fig. S4A). The bending frequency and the semiperiod SD did not show significant differences in these subsampled roaming states (Fig. S4B, C). The control animals’ trajectories were also spread in phase space (fig. S4E-G). Together, these data suggest that SAA plays an important role in modulating different aspects of bending kinematics, stabilizing the dynamics of coordinated rhythmic motion, and thus enhancing movement efficiency during roaming behavior.

**Fig. 3.**
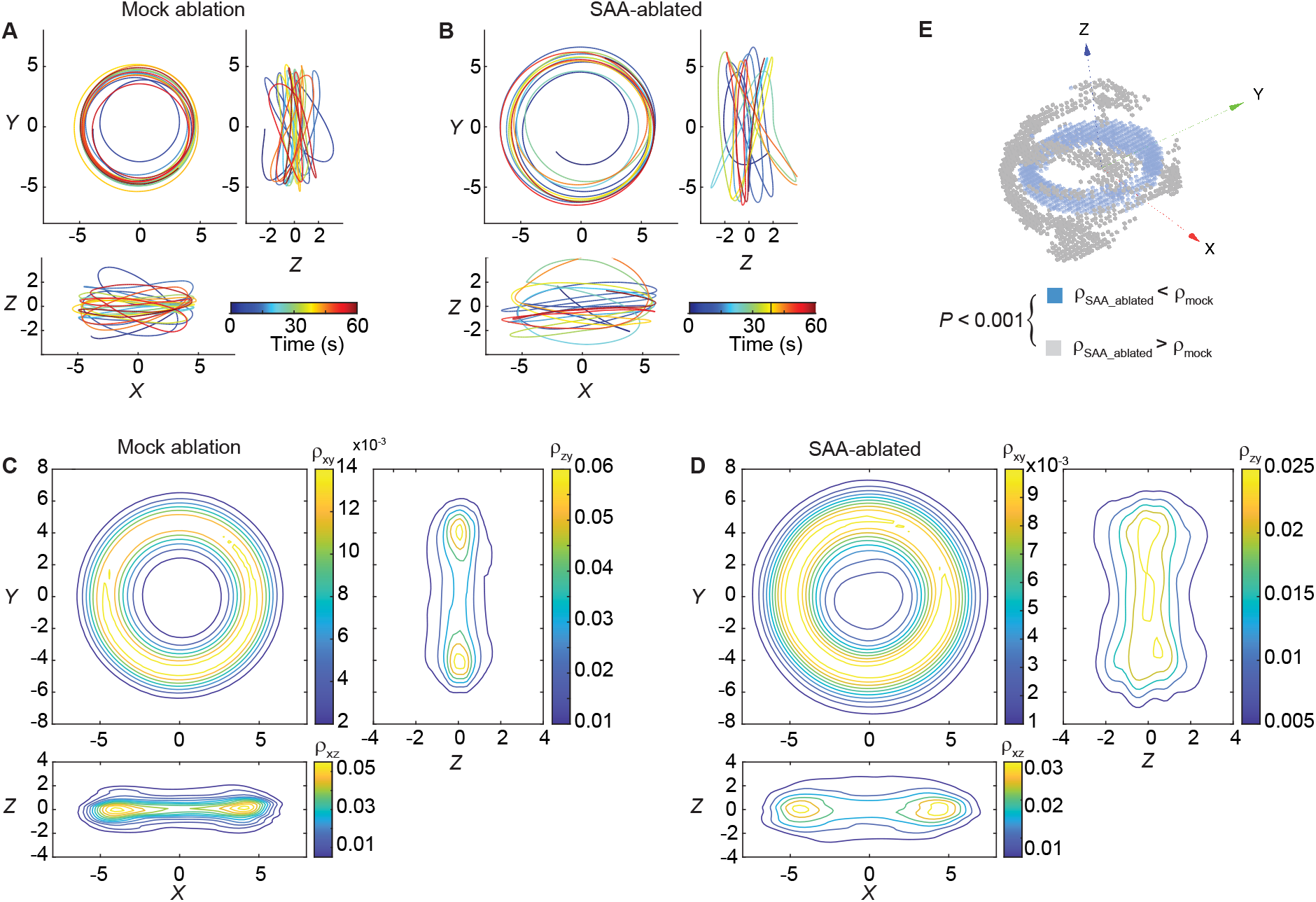
SAA stabilize rhythmic motion in forward movements. **A, B**. Time evolution of a phase trajectory in a control animal (**A**) and an SAA-ablated animal (**B**). Time is color-coded along the phase curves to represent the progression of movement. **C**. Density of trajectories embedded in the 3-dimensional phase space during forward movements of control animals. Each of the three subpanels represents a density projection onto a plane spanned by two orthogonal directions. **D**. Similar to **C**, but for SAA-ablated animals. **E**. Local density differences between **C** and **D** were visualized by a voxelgram (Methods). The experiments in this figure were carried out on 0.8% agarose pad without OP50 lawn. Ctrl: n = 113, 8 animals; SAA-ablated: n = 85, 8 animals.

The wiring diagram of the head motor circuit (fig. 2A) implies that SAA has the ability to assimilate and modulate the functioning of head motor neurons through a mixture of chemical and electrical synapses. Consequently, it is logical to ask the connection between the activity of SAA and complex head movements. fig. 2G presents typical ratiometric Ca^2+^ activity patterns of SAAD and SAAV during an extended recording session (140 s) that covers various behavioral states (Methods). As the worm moved forward, SAAD displayed rapid oscillatory behavior (fig. 2G and fig. S5) which is strongly correlated with the alteration of head curvature (fig. 2H). Consistent with this finding, the power spectrum density (PSD) of SAAD Ca^2+^ activity revealed a noticeable local peak close to 0.1 Hz that matched the highest power density of head bending activity (fig. 2I). However, we noticed a reduction in the correlation between Ca^2+^ activity and head bending activity during reversals (fig. 2H,I). Moreover, during the transition from reversal to turn, SAAD demonstrated increased Ca^2+^ activity (fig. 2G), an occurrence that we will examine later. Interestingly, on the short timescale, we did not observe robust state-dependent Ca^2+^ activity in the SAAV neuron (fig. 2H), suggesting a functional asymmetry that cannot be inferred directly from the anatomical connectome.

### Impact of SAA on the long timescale locomotory behavior

We monitored the crawling behavior of *C. elegans* on an empty agar pad after the animal was removed from food for approximately 15 minutes, which corresponds to the global dispersion state. On this long timescale, SAA-ablated animals made more frequent transitions to other motor states (fig. 4A,B). In fact, the percentage of recording time during which the animals spent in reversal, pause, and turn significantly increased (fig. 4B). The cumulative density function (cdf) curve of the duration of the forward run shifted upward in SAA-ablated animals (fig. 4C), suggesting a significantly shorter run duration (fig. 4D). In contrast, the duration of spontaneous reversal was longer in SAA-ablated animals (fig. 4E,F). A similar change in behavior was observed in swimming animals (fig. S6). Consistent with this observation, during optogenetic inhibition of SAA (P*lim-4*::loxP::Arch;P*Lad-2*::Cre), animals were more likely to make reversals (fig. 4G).

**Fig. 4.**
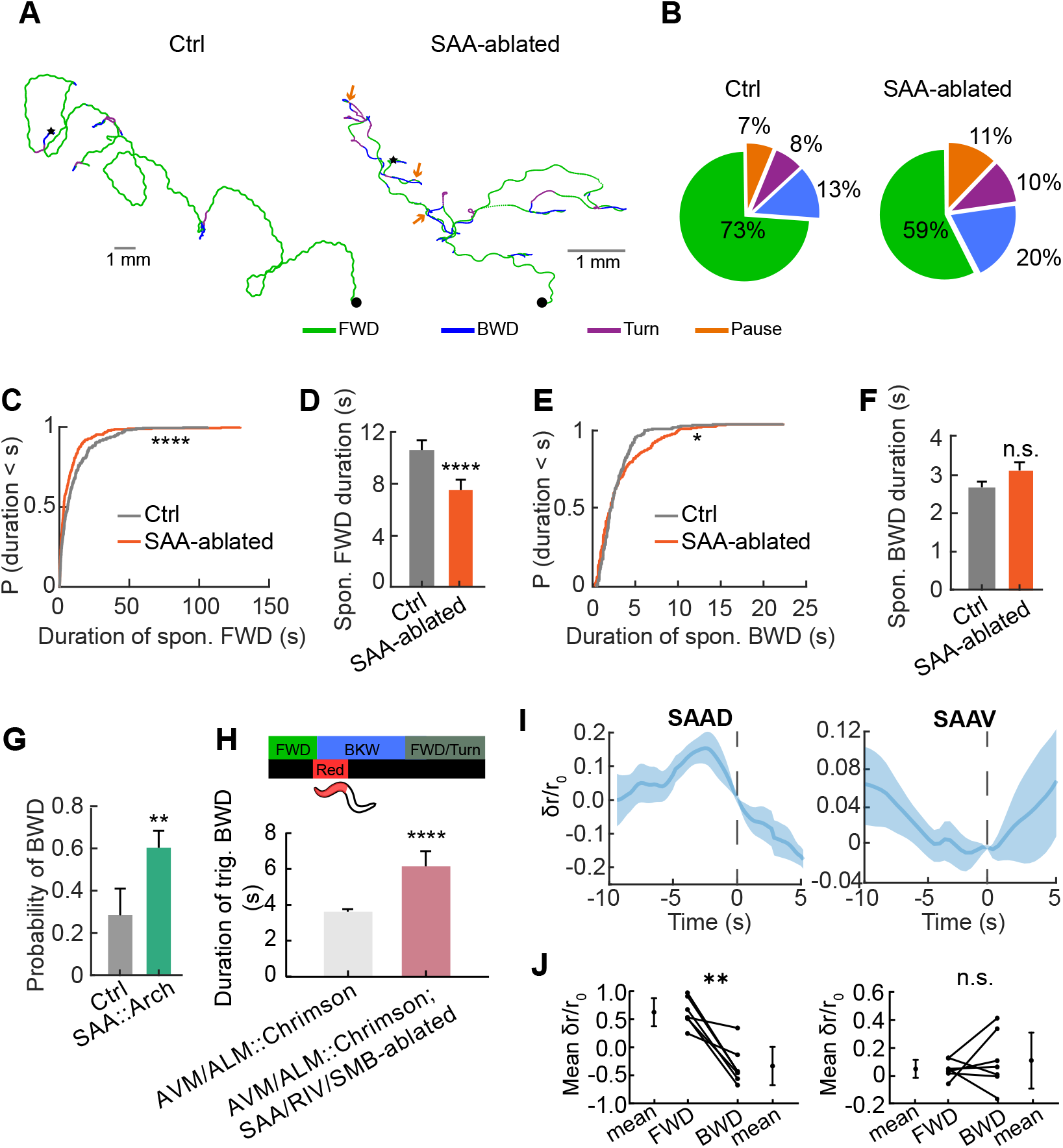
Impact of SAA on the long timescale crawling behaviors. **A**. Representative crawling trajectories of a control and a SAA-ablated animal. Different motor states are color-coded, and arrows indicate the pause state. Each worm was recorded for about 5 minutes. **B**. The percentage of time spent on forward movements, reversals, turns, or pauses. ****P *<* 0.0001, *χ*^2^ test was performed on fractional time spent in the reversal, turn, and pause states. The control group represents mock ablated animals. **C**. Cumulative distributions of the forward run length. Related to **B**. ****P *<* 0.0001, two-sample Kolmogorov–Smirnov (KS) test. Ctrl: n = 308, 8 animals; SAA-ablated: n = 273, 8 animals. **D**. Mean duration of spontaneous forward runs in control and SAA-ablated animals. ****P *<* 0.0001, Mann–Whitney U test. The error bars represent SEM. **E**. Cumulative distributions of reversal length. Related to **B**. *P = 0.048, two-sample KS test. Ctrl: n = 214, 8 animals; SAA-ablated: n = 220, 8 animals. **F**. Mean duration of spontaneous reversals in control and SAA-ablated animals. The error bars represent SEM. **G**. The probability of eliciting a spontaneous reversal when SAA interneurons were optogenetically inhibited for 7 s by a green laser. The interval between optogenetic manipulations was *>* 45 s. The control group was fed OP50 without all trans-retinal. **P *<* 0.01, *χ*^2^ test. Error bars indicate the 95% confidence interval for the binomial proportion. Ctrl: n = 39, 7 animals; SAA::Arch: n = 85, 18 animals. **H**. Top: illustration of the experimental procedure for inducing escape behavior by activating AVM/ALM (P*mec-4*::Chrimson). The anterior half of an animal body was illuminated by 1.5 sec red light during a forward movement. Bottom: duration of ALM/AVM-triggered reversals in control animals and SAA-ablated animals. ****P *<* 0.0001, Mann–Whitney U test. The error bars represent SEM. AVM/ALM::Chrimson: n = 414, 61 animals; AVM/ALM::Chrimson;SAA/RIV/SMB ablated: n = 41, 8 animals. **I**. Changes in normalized neuronal activity before and after reversal onset. The moment of reversal initiation is indicated by 0 s. The shaded region indicates SD. Trials were included if they had long forward movement (*>* 5 sec) followed by long reversal (≤ 5 sec). Displayed as *δ r/r*_0_ = (*r*(*t*) − *r*(*t* = 0))*/r*(*t* = 0). The shaded areas in the two curves represent SEM. n = 7, 7 animals. **J**. Related to **I**, average normalized neuronal activities for the 5-second interval preceding and following the initiation of reversals. **p = 0.007 for SAAD, n.s. p = 0.49 for SAAV, using a paired t-test.

How does the activity of SAA change during the forward-to-reversal transition? When aligning the Ca^2+^ activity with the start of a reversal (*t* = 0), we found that the SAAD activity showed a gradual decline prior to the transition (fig. 4I) and there was a notable difference in the average activity before and after the transition (fig. 4J). In contrast, the trial-averaged SAAV activity did not show distinct changes during the transition from forward movement to reversal (fig. 4I and fig. 4J).

SAA affected not only long-timescale spontaneous behaviors but also stimulus-triggered motor state transitions. Here, we quantitatively characterized escape responses in transgenic animals (P*mec-4*::Chrimson) induced by optogenetically activating mechanosensory neurons ALM/AVM (fig. 4H). Ablation of SAA (P*lim-4*::PH-miniSOG, PH-miniSOG was also expressed in RIV and SMB neurons) led to longer reversals, suggesting that the ability to terminate reversals via backward-turn transitions was impaired (35).

### Activation of SAA facilitates reversal termination

What is the neural basis underlying the long-timescale behavioral changes observed in SAA-ablated animals? According to the *C. elegans* connectome, SAA make prominent chemical synapses with several interneurons (including RIM/AVA/AIB) that control backward movements (fig. 5A). One possibility is that SAA activity can directly modulate motor state transitions through these interneurons. To test this hypothesis, we designed an experiment to optogenetically activate SAA during the reversal state. In order to avoid light spectra overlap, here we triggered an escape response by thermally stimulating the worm head for 1 second, followed by 7-second optogenetic activation of SAA (fig. 5B, Methods). We constructed transgenic animals in which Chrimson was expressed specifically in SAA (P*lim-4*::loxP::Chrimson;P*Lad-2*::Cre) or in SAA/RIV/SMB neurons (P*lim-4*::Chrimson). Activation of SAA/RIV/SMB or SAA alone could rapidly terminate reversals: *termination latency*, defined as the time between the onset of optogenetic stimulation and the end of a reversal, was significantly shorter than in control animals (fig. 5C and Methods). Furthermore, stimulation of SAA / RIV / SMB while blocking the chemical synaptic transmission from these neurons (P*lim-4*::TeTx) prolonged the latency (fig. 5D). SAA form numerous connections with SMB. To avoid confounding factors arising from signal propagation between SAA and SMB, we ablated SMB specifically using a Femtosecond laser (fig. 5E). Similar to fig. 5C, the triggered reversal was quickly terminated by SAA activation (fig. 5E). Consistent with the optogenetic experiment, we found that during the reversal-turn-forward transition, SAAD / SMB neurons exhibited elevated calcium activity (fig. 5F). Together, these results suggest that the feedback synaptic inputs from the depolarized SAA to the interneurons in the backward module facilitate the termination of the reversal.

**Fig. 5.**
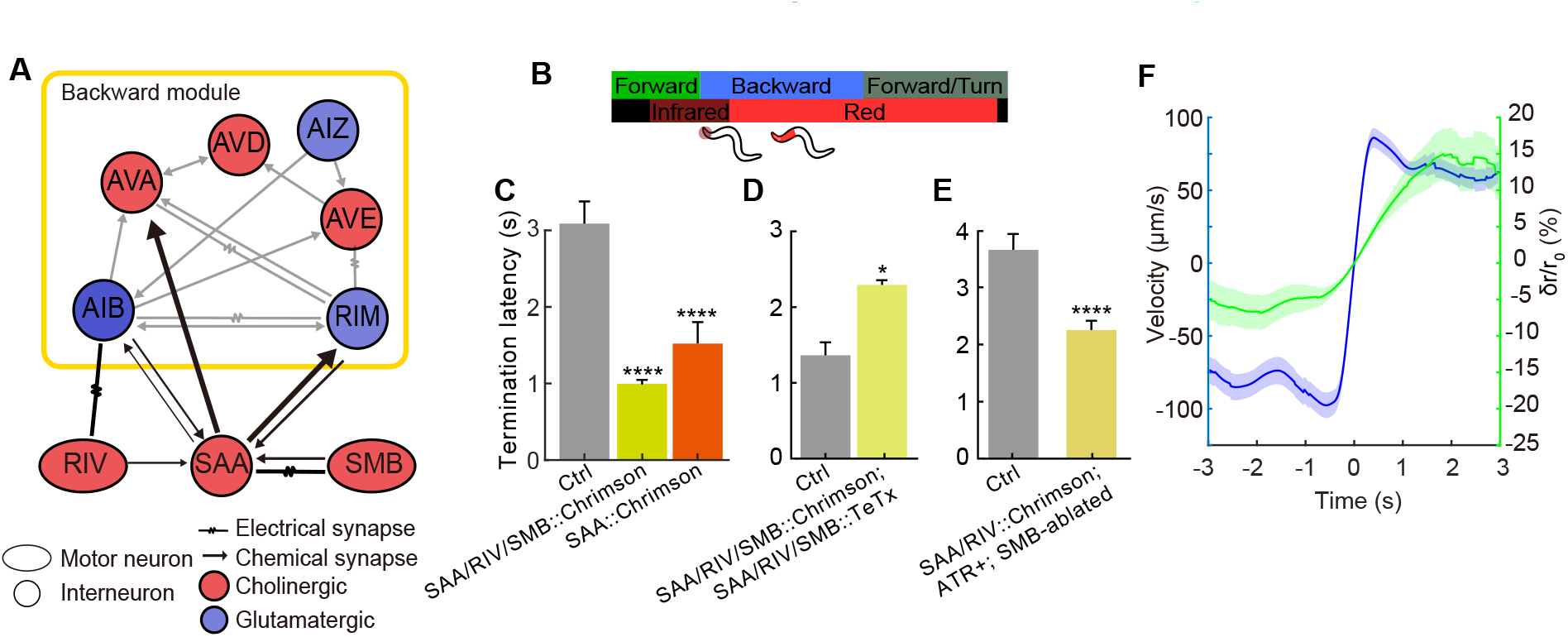
Activation of SAA facilitates reversal termination. **A**. The neuronal circuit bridging lower level head motor circuit and higher level command center. The synaptic convergence and divergence of SAA are proportional to the width of the line. For example, the number of synapses from SAA to RIM is *≈* 70 and the number of synapses from RIM to SAA is *≈* 30. **B**. Schematic experimental procedure for activation of SAA or SAA/RIV/SMB during thermally induced escape responses. Reversal was triggered by an infrared laser that focused on the head of the worm followed by optogenetic stimulation (red light). Related to **C**-**D. C**. Termination latency between the onset of optogenetic stimulation of SAA neurons or SAA/RIV/SMB neurons and the end of a reversal. ****P *<* 0.0001, compared to the control group, Mann-Whitney U test with Bonferroni correction. Ctrl: n = 52, 10 animals; SAA::Chrimson: n = 36, 9 animals; SAA/RIV/SMB::Chrimson: n = 229, 53 animals. **D**. Similar to **C**, but in one group, chemical synaptic transmission from SAA/RIV/SMB was blocked by an expression of tetanus toxin. *** P *<* 0.001, compared to the control group, Mann-Whitney U test. SAA/RIV/SMB::Chrimson: n = 65, 11 animals; SAA/RIV/SMB::Chrimson;SAA/RIV/SMB::TeTx: n = 60, 14 animals. **E**. Termination latency between the onset of optogenetic stimulation of SAA/RIV neurons with ablated SMB neurons and the end of a reversal. ****P *<* 0.0001, Mann–Whitney U test. Ctrl (SMB-ablated worms fed without ATR): n = 58, 6 animals; experiment group: n = 89, 13 animals. **F**. Calcium imaging of SAAD/SMB near the reversal-turn transition. t = 0 was aligned with the reversal end (that is, velocity = 0 mm/s). The blue curve is the mean velocity of worm movements; green curve is the mean ratiometric calcium signal in SAAD/SMB neurons, plotted as *δ r/r*_0_ = (*r*(*t*) − *r*(*t* = 0))*/r*(*t* = 0). The shaded areas in the two curves represent SEM. n = 40, 13 animals.

### Inhibitory acetylcholine synaptic transmission promotes the termination of reversals

SAA are cholinergic neurons (45), and *C. elegans* nervous system possesses a family of acetylcholine-gated chloride (ACC) channels, and several putative subunits have been identified, including ACC-1 to ACC-4 (45, 46). Therefore, we first ask whether these inhibitory acetylcholine receptors are involved in motor state transitions. We crossed transgenic animals (P*mec-4*::Chrimson) expressing Chrimson in mechanosensory neurons with ACC-deficient mutants: *acc-1(tm3268), acc-2(tm3219), acc-2(ok2216), acc-3(tm3174), acc-4(ok2371)*. Optogenetic activation of ALM and AVM were able to trigger prolonged reversals in these mutants (fig. 6B). Interestingly, in two of the double mutants we tested (*acc-2(tm3219)*;*acc-3(tm3174)*, and *acc-3(tm3174)*;*acc-4(ok2371)*), the reversal duration was significantly longer than that of a single mutant (fig. S7), indicating that these channel subunits act synergistically in the nervous system. Likewise, using the thermal and optogenetic stimulation protocol (Methods), we found that the termination latency increased in ACC-deficient mutants (fig. 6I and fig. S8B-D).

**Fig. 6.**
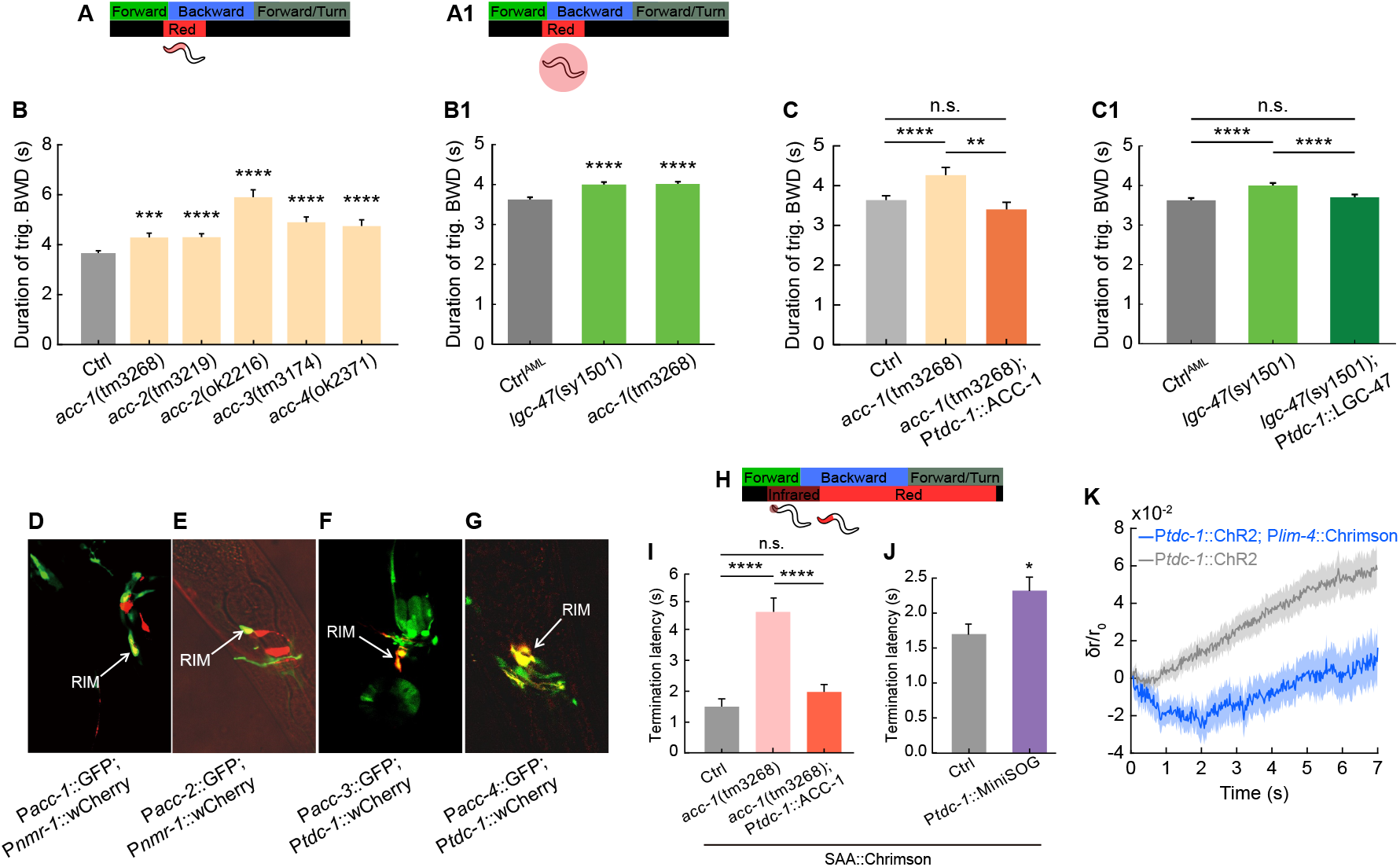
RIM communicates with SAA to terminate reversals via inhibitory cholinergic synapses. **A**. Schematic experimental procedure for triggering escape responses (same as fig. 4H). Optogenetic stimulation would activate AVM/ALM mechanosensory neurons. Related to **B** and **C. A1**. Schematic experimental procedure for triggering escape responses. Illumination of the entire worm activates all six touch receptor neurons (ALM L/R, AVM, PLM L/R, PVM). Related to **B1** and **C1. B**. Duration of ALM/AVM-triggered reversals in N2 (wild type) and ACC-deficient animals. Ctrl: n = 414, 61 animals; *acc-1(tm3268)*: n = 114, 16 animals; *acc-2(tm3219)*: n = 100, 10 animals; *acc-2(ok2216)*: n = 88, 12 animals; *acc-3(tm3174)*: n = 97, 11 animals; *acc-4(ok2371)*: n = 123, 16 animals. **B1**. Duration of all-TRN-triggered reversals in Ctrl, LGC-47 deficient, and ACC-1 deficient animals. Here a different mec-4 Chrimson allele is used, denoted by AML. The number of optogenetic stimulus events from left to right is 550, 498, and 624. **C**. Duration of ALM/AVM-triggered reversal in N2, *acc-1* mutant, as well as animals in which ACC-1 was specifically restored in RIM. Ctrl: n = 414, 61 animals; *acc-1(tm3268)*: n = 114, 16 animals; *acc-1(tm3268)*;P*tdc-1::ACC-1*: n = 101, 16 animals. **C1**. Duration of all-TRN-triggered reversal in Ctrl, lgc-47 mutant, as well as animals in which LGC-47 was specifically restored in RIM. The number of optogenetic stimulus events from left to right is 550, 498, and 341. **D-G**. GFP reporter lines show a co-localization of acc-1, acc-2, acc-3, and acc-4 with RIM interneuron. **H**. Schematic procedure for dual thermal and optogenetic stimulation (same as fig. 5B). Optogenetic stimulation would activate SAA neurons. Related to **I**-**J. I**. Termination latency in control, *acc-1* mutant, as well as animals in which ACC-1 was specifically restored in RIM. Ctrl: n = 35, 8 animals; *acc-1(tm3268)*: n = 48, 10 animals; *acc-1(tm3268)*,P*tdc-1::ACC-1*: n = 84, 15 animals. **J**. Termination latency in control and RIM ablated animals. Ctrl (mock-ablated): n = 145, 26 animals; RIM-ablated: n = 64, 13 animals. In the experiments detailed in (**I**) and (**J**), SAA neurons were selectively activated. All Statistical tests: *P=0.010, **P *<* 0.01, ****P *<* 0.0001, Mann–Whitney U test or Mann–Whitney U test with Bonferroni correction. Error bars represent SEM. **K**. Calcium dynamics of RIM neurons. Upon blue light excitation at t = 0, the GCaMP signal was monitored in RIM. The co-expression of ChR2 in RIM would simultaneously depolarize the neuron during imaging, leading to a continuous increase in the calcium signal (gray). Activation of RIM together with SAA/SMB/RIV led to a transient but significant decrease in calcium activity in RIM (blue) shortly after stimulation onset. The animals were immobilized on an 10% agarose pad with a coverlip. P*tdc-1*::ChR2: n = 39, 7 animals; P*tdc-1*::ChR2;P*lim-4*::Chrimson: n = 21, 4 animals. Lines and shaded areas represent the mean *±* SEM.

### RIM communicates with SAA to terminate reversals

Acetylcholine-gated chloride channels have a wide distribution within the *C. elegans* nervous system. Which neurons receive synaptic input from SAA to terminate reversal? Using GFP reporter lines, we focus on overlaps of acc expression patterns with neurons known to encode backward motor states (fig. 5A). We found that, intriguingly, all four (*P*acc-1 to *P*acc-4) reporters exhibited expression in RIM (fig. 6D-G), an integrating interneuron that plays a central role in motor state transitions (35, 36, 47, 48). A recent study (36) demonstrated that a depolarized RIM promotes spontaneous reversal, while a hyperpolarized RIM suppresses reversal (also see fig. S8F,J) by exploiting a combination of chemical and electrical synapses (fig. 5A). The presence of four different types of acetylcholine-gated chloride channel subunits may coordinate to regulate RIM activity. Consistent with this notion, after restoring the expression of each of the 4 ACC channels specifically in RIM, we found that restoring ACC expression in RIM significantly reduced the duration of reversals triggered by ALM/AVM activation (fig. 6C and fig. S8G-I).

It has recently been proposed that ACC-1 may form a heterameric ion channel with LGC-47(49, 50). We therefore tested the role of LGC-47 on reversal duration length using a separate set of strains. The defects in LGC-47 resulted in prolonged reversal duration similar to those of ACC-1 defective animals (fig. 6B1). Restoring LGC-47 only in RIM reduced the duration of the reversals (fig. 6C1), while restoring LGC-47 in other interneurons did not (fig. S8K). Together, this suggests that LGC-47 in RIM is also part of the machinery that receives inhibitory cholinergic signals from SAA to modulate reversal duration.

We further explored how synaptic input from SAA to RIM would shape the timescale of reversals. Using the thermal and optogenetic stimulation protocol, we found that restoring ACC subunit expression specifically in RIM significantly reduced termination latency (fig. 6I and fig. S8B and C). In RIM-ablated animals, the termination latency, on the other hand, significantly increased (fig. 6J). Note that these experiments (fig. 6I,J) were carried out by selectively activating SAA during a reversal. Finally, we directly observed a significant decrease in RIM calcium activity (P*tdc-1*:GCaMP6; P*tdc-1*:ChR2) immediately after optogenetic activation of SAA. As a control, we co-expressed channelrhodopsin (ChR2) in RIM to ensure that the neuron would be in a depolarized state during calcium imaging (fig. 6K). Together, these experiments suggest that cholinergic synaptic inhibition from SAA to RIM contributes to the termination of the reversal and promotes forward movements.

## Discussion

### Summary

The neural basis of animal behavior, ranging from fast body movements to slow exploration-exploitation strategy, has been extensively studied in the literature (12, 13, 21, 32, 51– 55). Here, we show that in *C. elegans*, the interneurons SAA, which are located in the lower level head motor circuit, bridge the short time-scale dynamics that characterize rhythmic bending activity and the long time-scale dynamics that describe motor state transitions (fig. 7).

**Fig. 7.**
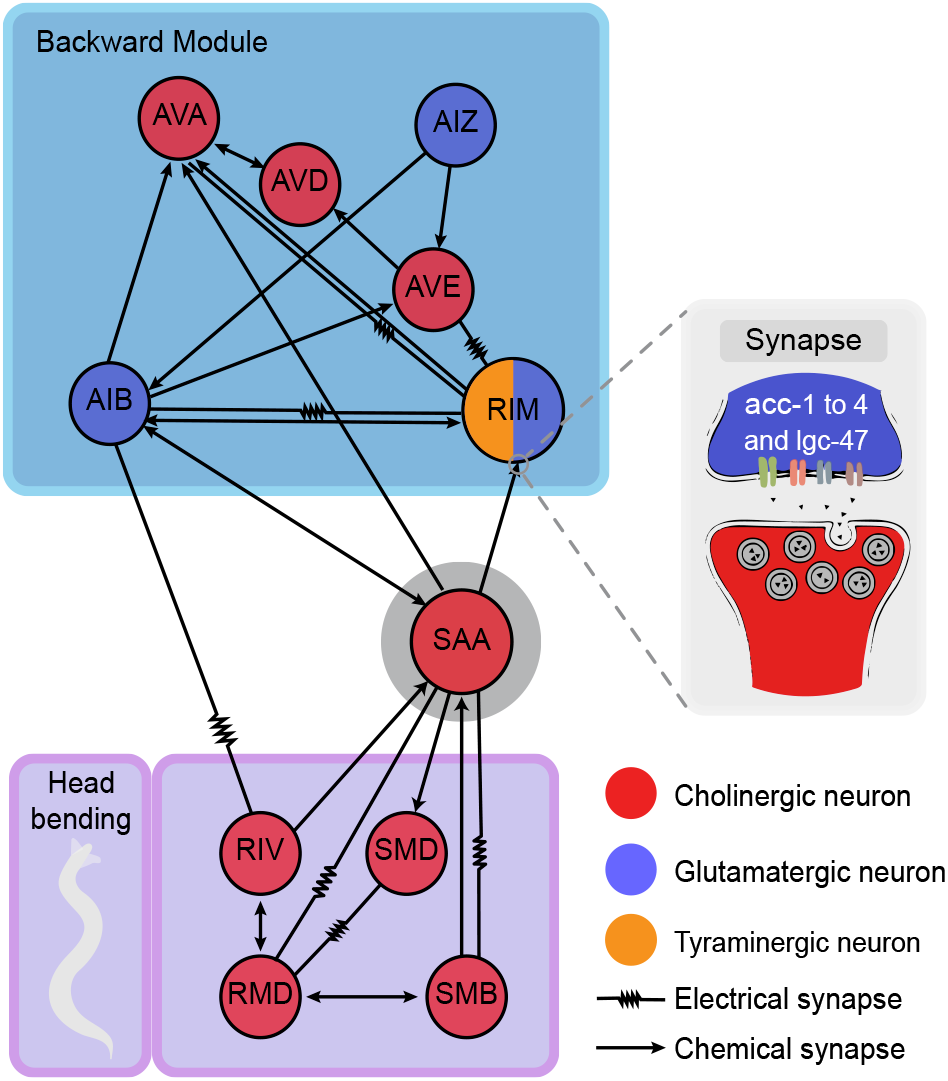
SAA interneurons bridge circuit modules that control short and long timescale motor behaviors.

On the short timescale, we find that SAA interneurons modulate the fast bending kinematics during forward movements, such as the undulation frequency and amplitude (fig. 2). The disruption of movement kinematics reduces locomotion speed (fig. 2, fig. S2 and S4). In particular, the impact of SAA is not restricted to head movements; instead, the modulation is translated into the whole body of the worm (fig. 3). SAA do not form neuromuscular junctions directly, but they make numerous chemical synapses and gap junctions with cholinergic head motor neurons, including SMB, SMD and RMD (fig. 2A). In fact, the exaggerated amplitude of body undulation (fig. 2F) in SAA-ablated animals was similarly observed in SMB-ablated animals (24). How SAA interneurons mediate activities in these motor neurons remains to be identified. The worm connectome (fig. 2A) indicates that SAA facilitate cross-coupling between dorsal and ventral head motor neurons. This bilateral cross-coupled motif could be essential for stabilizing rhythmic motion by setting the correct phase difference between dorsal and ventral motor activities. In addition, SAA have processes that extend anteriorly without presynaptic specializations, a feature that led White (41) to hypothesize that these processes have proprioceptive properties. A candidate for a potential mechanosensory channel is the TRP family (56), and TRP1 is known to express in SAA (57). Previous works (28, 30, 58, 59) suggest that when swimming in environments with different mechanical loads, *C. elegans* exhibited gait adaptation by exploiting local proprioceptive feedback. Consistent with this view, SAA-ablated animals exhibited a large variation in the undulatory period when swimming in solutions with changing viscosities (fig. S2C-D). Whether this observation results directly from a loss of proprioceptive signal, an impairment of the cross-coupled circuit, or both remains to be understood.

On the long timescale, we find that SAA interneurons modulate motor state transitions by providing feedback inhibition to RIM, an integrating neuron that modulates the frequency and duration of a reversal (36, 47, 48, 60). A bidirectional change in RIM activity can promote or suppress the reversal state (36, 48). During stimulus-triggered escape responses, feedback inhibition is part of the winner-take-all strategy (35) that ensures a rapid and smooth transition from a reversal state to a turn/forward movement state, accompanied by increased calcium activity in SAAD (fig. 5F). This inhibition is achieved by cholinergic synapses. Interestingly, we find that RIM expresses all four acetylcholine-gated chloride channel subunits, and our data suggest that different ACC channel subunits do not act redundantly (fig. S7). Here, we postulate that a spatial combination of ACC subunits in the RIM cell body or neurites may contribute to fine-tuning the RIM membrane potential and enable precise control of motor state transitions.

### Functional diversity of interneurons

In *C. elegans*, interneurons situated between sensory and motor neurons play crucial roles in action selection and modulation of adaptive motor behaviors. Interneurons such as AIY and AIB, which receive substantial sensory inputs, demonstrate a rapid transition from sensory to motor representation: Their Ca^2+^ activities encode distinct motor states over extended timescales. Disrupting these neurons alters the likelihood of transitions between motor states without affecting rapid fast motor kinetics, such as movement speed (fig. S10) (27, 35, 61). Descending the sensorimotor hierarchy, interneurons with extensive connections to motor neurons, such as AVB, AVA, and RIB, show a different function. Manipulation of these neurons through optogenetic stimulation or inhibition can initiate or stop movements, respectively (26, 30, 35, 37). Their removal or persistent inhibition of these neurons significantly reduces movement speed (fig. S10 and (37)) and the amplitude of undulation (30), indicating that sustained activity in AVB, AVA and RIB (26, 27, 31, 33, 35, 37) provides a feedforward gating signal that facilitates forward movements and reversals. These three gating neurons maintain movement states rather than finetuning kinematics on the timescale of an undulation period. The decreased movement speed and undulation amplitude are due to the lack of sustained activation signals to motor neurons, but their calcium activity does not correlate with fast undulation activity (for example, see RIB Ca^2+^ activity in fig. S12) and (26, 30, 35, 37).

SAA interneurons are different from most interneurons in that their calcium activity reflects both fast dynamics correlated with head-bending and slower state-dependent fluctuations (fig. 2G). Our experiments with optogenetics and neuron ablation reveal that SAA serves a dual function: it stabilizes rapid undulatory dynamics and modulates transitions between motor states, thus complementing the role of command interneurons. This finding, together with previously described SMD motor neurons (13), introduces a class of vertically integrated neurons that traverse different timescales. It should be mentioned that RIA interneurons, which receive an efferent copy from SMD motor neurons, also display fast activity associated with head bending during navigation (62, 63). However, despite influencing the direction of head movements, RIA appears to be not directly involved in action selection or the control of movement kinematics (fig. S11) and (24).

#### Feedback loop in the motor control hierarchy

The concept of behavior hierarchy (fig. 1) provides a compelling framework to organize the intricate dynamics of complex systems. Various studies of the *C. elegans* motor circuit now suggest that feedback loops exist at each hierarchical level. At the bottom level, proprioceptive feedback within motor neurons dictates the frequency and wavelength of undulatory movements (28, 30, 39). At the intermediate level, we observe that the activity state of motor neurons can influence action selection retrogradely. For example, decreased activity in head motor neurons SMD is correlated with prolonged periods of reversal (13). This observation is consistent with the function of SAA: When head motor activity is subdued, there is insufficient feedback inhibition to rapidly terminate the reversal. As we move further along the hierarchy, the command neurons responsible for reversals, namely RIM and AVA, form intricate recurrent connections with interneurons such as AIB and AIY, where sensory and motor state information are integrated (47, 48, 64, 65).

What computational advantages does feedback inhibition provide when it occurs from the lower-level motor circuit to the higher-level control center? We believe that this strategy could efficiently stabilize behavioral states. During navigation, RIM and other interneurons within the backward module (fig. 5A) are bombarded with noisy sensory input. Ongoing synaptic inhibition from SAA would hyperpolarize RIM, thus helping maintain a global dispersion state (fig. 4). Consistent with this notion, we observed that inhibition of SAA increases the frequency of reversals (fig. 4G); similarly, animals with ablated RIM exhibited more frequent reversal transitions (fig. S9). Bottom-up feedback connections (fig. 1) therefore ensure robust behavioral control by complementing a rigid top-down hierarchy.

This organizing principle is broadly relevant in vertebrate and mammalian systems. For example, gap junction-mediated retrograde signal propagation from motor neurons to interneurons has also been reported in the spinal cord of zebrafish (66). In mammalian systems, complex feedback structures such as corticospinal feedback, cerebellar-spinal feedback, and the Cortico-Basal Ganglia-Thalamo-Cortical loop exemplify the sophisticated architecture of the motor control hierarchy (67– 72). These feedback loops play a crucial role in regulating movement decisions, facilitating motor learning, and ensuring precision and smoothness of movements through ongoing error correction. The study of *C. elegans* therefore provides a window into the fundamental algorithms that govern adaptive motor behaviors. The difference lies in the degree of specialization: Whereas the compact nervous system of *C. elegans* may use a single, multifunctional neuron type to control behavior, mammalian systems often distribute these functions between specialized cell types and dedicated brain regions to manage more complex behaviors.

## Materials and Methods

### Strains

Wild-type (N2), mutants, and transgenic animals were cultivated using standard methods. The specific promoter-driven expression of Chrimson, Arch, GCaMP6, miniSOG, or Tetanus toxin were co-injected with P*lin-44*::GFP or P*unc-122*::GFP injection marker into N2 to generate transgenic animals. P*mec-4*::Chrimson (WEN1015) and P*lim-4*::Chrimson (WEN1009) were integrated by a UV illumination method and outcrossed 6X with N2 animals. The integrated lines were crossed with *acc-1, acc-2, acc-3*, and *acc-4* mutants. To manipulate SAA selectively, we used Cre*/*loxP system to generate transgenic animals that Chrimson, Arch and miniSOG were expressed in SAA neurons. P*lad-2*::Cre and P*lim-4*::loxP::Chrimson*/lim-4*::loxP::Arch*/lim-4*::loxP::PH-miniSOG were mixed and co-injected with P*lin-44*::mCherry injection marker into N2. The transgenic animals used in all optogenetic experiments were grown in the dark at 16 to 22 °C on NGM plates with *Escherichia coli* OP50 and all-trans retinal (ATR) for over 8 hours. All experiments were carried out with young adult hermaphrodites. Detailed information can be found in the Supplemental Tables (S2).

#### Molecular biology

Standard molecular biology methods were used. Promoters such as P*lad-2*(2.1 kb), P*acc-1*(4.0 kb), P*acc-2*(2.0 kb), P*acc-3*(4.0 kb), and P*acc-4*(1.5 kb) were amplified by PCR from the genome of wild-type animals. Detailed information can be found in the Supplemental Tables (S2).

#### Behavior recording

*C. elegans* were placed on a 0.8% (wt/vol) M9 agar plate or immersed in dextran solution (5, 15, 25 % (wt/wt) dextran in M9 buffer). Before recording, the worms were transferred to a sterile NGM plate to remove bacteria from the body surface and then transferred to the recording device: an agarose plate (0.8% (wt/vol) agarose in M9 buffer) or a chamber with dextran solution. The animals acclimatized to the new environment for 10 to 15 minutes before being recorded freely for 5 to 10 minutes on a Nikon inverted microscope (Ti-U) with dark-field illumination.

#### Optogenetics

Experiments were performed on an inverted microscope (Ti-U, Nikon, Japan) at 10x magnification with dark-field illumination. The animals were placed on a 0.8% (wt/vol) M9 plate and retained within the field of view of an imaging objective using a custom tracking system. The video sequences were captured by a Basler CMOS camera (aca 2000-340km), and the worm body centerline was extracted in real time. We used MATLAB custom software (MathWorks, Natick, USA) for post-processing behavioral data. We used the CoLBeRT system (73) to perform spatiotemporal optogenetic manipulation. For optogenetic activation of ALM/AVM neurons, we used a 635-nm solid-state laser with an intensity of 4.6 mW/cm^2^; in each trial, the illumination lasted 1.5 s.

#### Whole worm optogenetic stimulation

Reversal durations in response to whole worm optogenetic stimulations reported in fig. 6B1, fig. 6C1, and fig. S8K only are new analyses of behavior measurements first reported in (74). That work used a high-throughput optogenetic delivery system to illuminate the entire worm and activate all six gentle touch neurons. A 1.5 mm diameter circle of 80 *μ*W/mm^2^ intensity of 630 nm light was centered on the worm for 3 s during each stimulation. Additional methods are described in (74).

#### Optogenetic ablation

Optogenetic ablation was performed using transgenic strains, in which miniSOG was expressed in target neurons. We used mitochondrially targeted miniSOG (TOMM20-miniSOG) and membrane-targeted miniSOG (PH-miniSOG) to induce cell death under blue-light illumination. Well-fed L3/L4 animals were transferred to an unseeded NGM plate and their movements were restricted within an area using a piece of filter paper with a hole in the center. The diameter of the restricted area was 0.5 cm and the filter paper was soaked with 100 *μ*M CuCl^2^. The animals were illuminated for 15 minutes (PH-miniSOG) or 30 minutes (TOMM20-miniSOG) with blue light (M470L3-C5; Thorlabs) with an intensity of 133 mW/cm^2^. The temporal sequence consists of 0.5/1.5 s on/off pulses. The animals were recovered and grown to the young adult stage in the dark at 16 to 22 °C on NGM plates with *Escherichia coli* OP50. This long overnight recovery ensures that the target neurons were killed.

#### Thermally-induced escape responses combined with optogenetic stimulation

To trigger an escape response, we used a thermal stimulus by illuminating the head of a worm with a focused infrared laser (1480 nm, 5 mW/mm^2^). Animals responded with reversals to avoid the thermal stimulus. We used custom-written LabVIEW scripts (National Instruments, USA) to control the diaphragm shutters (GCI 7102M, Daheng Optics, China) along the optical path to achieve sequential light activation. We used the CoLBeRT system to perform spatially selective optogenetic manipulation for different neurons. The illuminated area is the anterior 30 percent of the body. 1 s infrared light stimulation (to trigger escape response) was followed by red (635 nm, 8.4 mW/cm^2^) or green light (561 nm, 16 mW/mm^2^) with a duration of 7 s to activate SAA/RIV/SMB neurons or inhibit RIM interneuron.To selectively activate SAA interneurons, 5.71 mW/mm^2^ laser intensity was used.

#### Calcium imaging

Calcium imaging was carried out in animals expressing GCaMP6s or wNEMOs (43) and wCherry in the same neurons. Calcium dynamics was calculated as a ratiometric change. We imaged neurons in animals crawling on a 2% (wt/vol) NGM agarose plate with a coverlide. In fig. 2 and fig. 4, the worms crawled freely without a coverlide. In fig. 5E during a reversal-turn-forward transition. Reversals were triggered by an infrared laser (during forward movement. To image calcium activity in RIM while simultaneously activating SAA (fig. 6K), we synchronized blue light (488 nm) and green light (561 nm) excitation. The worms were immobilized with a high concentration agarose pad [10% (wt/vol)]. The illumination lasted for 7 s and the inter-activation interval was *>* 45 s. The green and red light emission signals were collected using a Nikon Plan Apo 10X objective and separated using an optical splitter device (OptoSplit II, Cairn-Research, UK), each of which was then projected onto one half of an sCMOS sensor (Zyla 4.2, Andor, UK). Neurons of interest were automatically identified using custom-written MATLAB scripts. Due to the wider emission spectrum of wNEMO, which caused fluorescence bleedthrough into the reference (wCherry) channel, we made a correction to the activity ratio using the equation:

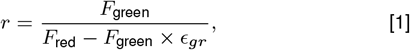

where *F*_*green*_ represents the fluorescence intensity in the green channel, *F*^*red*^ denotes the fluorescence value in the reference channel, *ϵ*^*gr*^ is the bleed-through ratio of wNEMO, and we have determined *ϵ* _*gr*_ to be 0.20.

#### Statistical test

All statistical tests were described in the figure legends, including methods, error bars, number of trials and animals, as well as p values. We applied the Mann-Whitney U test, the *χ*^2^ test to determine the significance of the difference between the groups, and the Kolmogorov-Smirnov test to compare the probability distributions. All multiple comparisons were adjusted using the Bonferroni correction. We performed the *χ*^2^ test using Excel and all other analyses using MATLAB.

#### Kinematic analysis of locomotion

Recorded videos and the corresponding yaml files were first processed to identify motor states (forward run, reversal, pause and turn). The timestamp, stage position, and the centerline of a worm were extracted using custom-written MATLAB scripts. To extract the bending curvature of an animal shown in fig. 2, the centerline of the worm was first divided into *N* = 100 segments and the orientation of each segment, θ (*s*),*s* = 1, 2,…, *N*, was computed. The curvature *κ* (*s*) = ∆ θ (*s*)*/* ∆*s* was calculated and then normalized into a dimensionless unit *κ* · *L*, where *L* is the body length. Spatial-temporal filtering was performed to obtain a smoothed version of *κ* (*s, t*) for further analysis.

To quantify the whole body bending amplitude, we first calculated the SD of curvature in a body segment

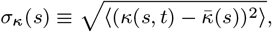

and we then averaged it on all segments 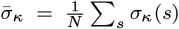. We smoothed the local curvature at a middle body segment and used the local peak finding algorithm in MATLAB to identify local maxima/minima of curvature in a time sequence. Semi-period is the time difference between adjacent peak and trough.

#### Phase space reconstruction and analysis

To reduce the spatiotemporal bending activity of worm movements to a trajectory in a low-dimensional phase space, we follow the procedure introduced in (44). First, by performing a principal component analysis on the worm postures, we approximated the intrinsic coordinate of each body segment, represented by a time-evolving *N* -dimensional curvature vector 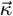, as a weighted sum of its leading principal components **e**^*i*^ (or eigenworms):

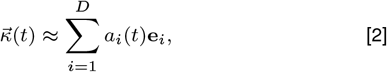

where the coefficients 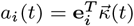, and we chose *D* = 5 such that the eigenworms captured more than 90% of the variance in worm posture data. Note that the eigenworm modes {**e**^*i*^} are consistent in the control and SAA-ablated datasets.

Next, we constructed a delay embedding matrix using the *D*-dimensional time coefficients (row vector) **a**(*t*) = [*a*^1^,…, *a*^*D*^]:

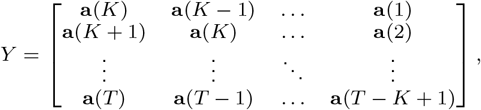

where *T* is the number of time frames in a single trial (forward run), *K* is the history time window. *Y* thus is a *T − K* +1 *KD* matrix. Here we chose *K* to be half of the mean undulation period (44), and typically *T ≫ KD*.

Finally, we performed singular value decomposition (SVD) of the *Y* matrix,

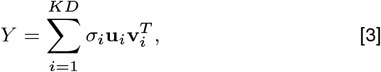

where *σ*^*i*^ is the singular value in descending order and **v**^*i*^ is the corresponding *KD*-dimensional basis vector. By examining the covariance matrix *Y* ^*T*^ *Y* with eigenvalues 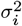, we found that with an embedding dimension *d* = 3, we could capture 94.1% variance in control animals and 89.6% variance in SAA-ablated animals during forward movements. Therefore, we projected *Y* onto the first 3 basis vectors, namely *X*^*i*^ = *Y* **v**^*i*^, *i* = 1, 2, 3, where *X*^*i*^ is a *T − K* + 1 column vector. The basis **v**^*i*^ was derived using three different approaches. (1) Each trial (forward run) independently generated a basis using a unique *Y* matrix. (2) We computed a consistent basis **v**^*i*^ for each animal by segmenting time sequences **a**(*t*) from each trial and merging them into a continuous sequence for each animal. (3) A consistent basis was applied for different animals. All three techniques produced comparable statistical results, specifically an expansion of the trajectories in the phase space (fig. 3D). Due to the behavior variability among animals, the direction that captures the most variance in the phase trajectory shows greater variation across animals than within trials of the same animal. Consequently, for visual representation, we opted for the second method in presenting our findings in fig. 3.

The spatial distribution of the phase trajectories was analyzed using the kernel density method in MATLAB and shown in a contour plot (fig. 3A,B). To compare the local density difference between control and SAA-ablated animals, we adopted an approach introduced in (75, 76), and used the kde.local.test function from the ks package in R to run a statistical test. Briefly, the phase space was first discretized into a 50 × 50 × 50 binning grid. Second, the local kernel density at a grid point was estimated, and the 3D spatial density function was represented by a kernel density matrix. Finally, pairwise local density comparisons between two matrices were performed with multiple comparison adjustment. Grid points that exhibited statistically significant (*P <* 0.001) differences were shown in fig. 3C.

## Supporting information

### Supplementary tables and figures

Supporting information comprises 3 supplementary tables containing information on plasmids and strains, as well as 9 supplementary figures.

## ACKNOWLEDGMENTS

We thank Zezhen Wang and Pinjie Li for helpful suggestions and preliminary analysis on the whole body bending kinematics. We thank Yixuan Li for the analysis of movement speed in the interneuron loss-of-function study. JH was supported by the National Natural Science Foundation of China, grant No. 82301666. TX was supported by the Young Scientists Fund of the National Natural Science Foundation of China, grant No. 32300829 and QW was supported by the Major International (Regional) Joint Research Project (32020103007)

## Supporting Information

### 1. Supplementary tables and figures

#### Supplementary figures (S1)

**Fig. S1.**
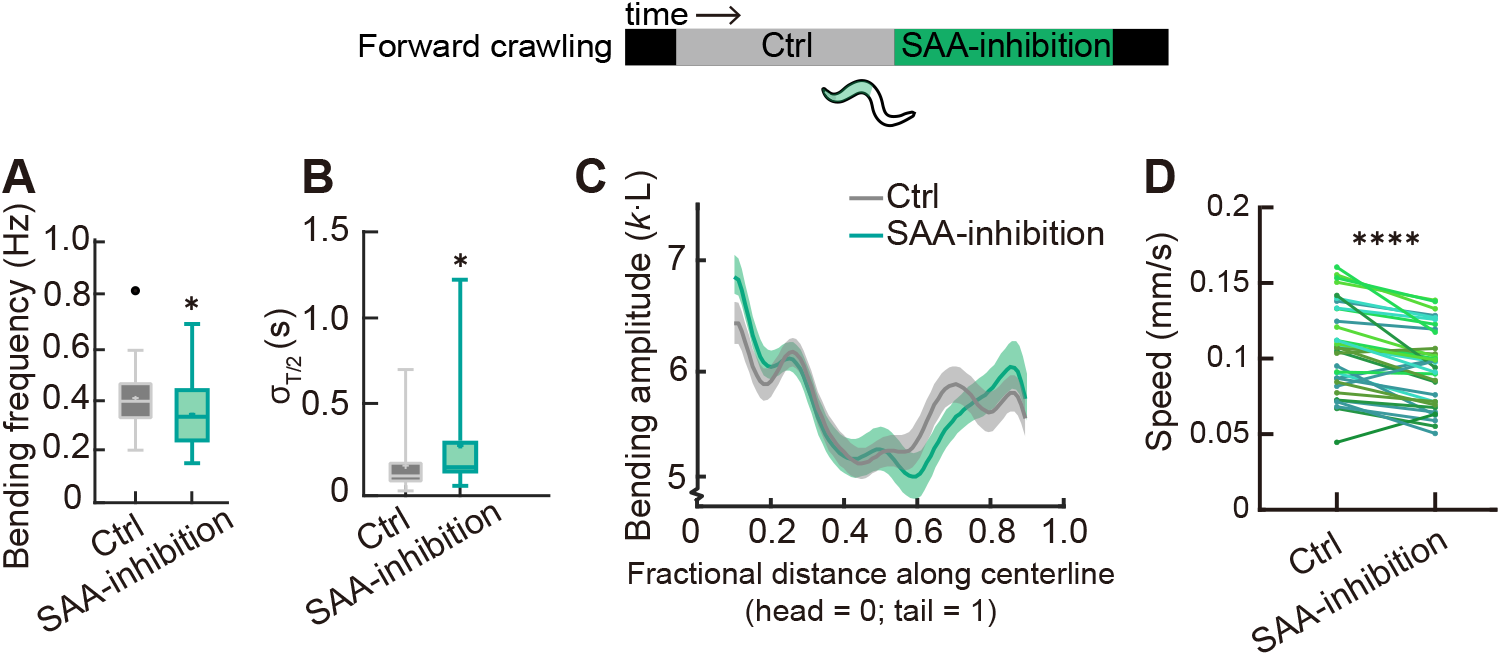
Acute inhibition of SAA affects fast kinematics in forward locomotion. **A**-**D**. Bending frequency, standard deviation of semi-period, bending amplitude as well as crawling velocity before and during SAA inhibiting within forward locomotion on agarose pad. *P = 0.0172 in **A**, *P = 0.0231 in **B**, ****P *<* 0.0001 in **D**, paired t-test. Box plots show the quartiles of the dataset, with whiskers extending from minimum to maximum, black dots are outliers and cross signs are mean. Solid curves are mean and shadow areas are S.E.M. Worms were illuminated by 561 nm laser to inhibit SAA. n=33, 10 animals

**Fig. S2.**
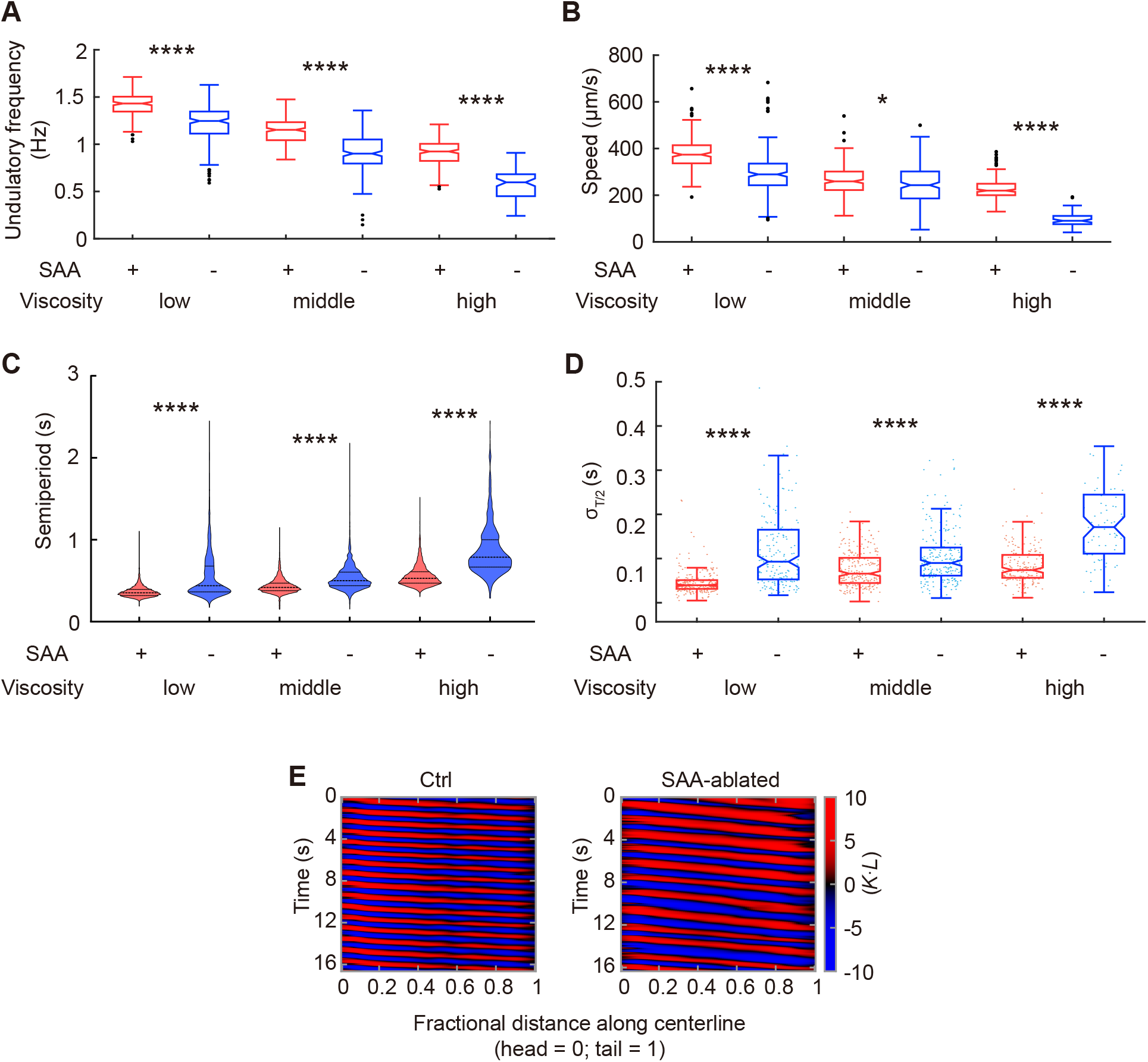
SAA contribute to stabilizing swimming kinematics. **A**-**C**. Velocity, undulation frequency, semi-period in control and SAA-ablated animals during forward locomotion in viscous solutions. **D**. Standard deviation of semi-period in control and SAA-ablated animals. Red and blue dots are standard deviations of each semiperiod of each trial. **A**-**D**: low, middle, or high viscosity represents 5% (9 mPa·s), 15 % (120 mPa·s), and 25% (800 mPa·s) dextran in M9 solution, respectively. *P *<* 0.05, ****P *<* 0.0001, Mann–Whitney U test with Bonferroni correction. Black dots are outliers. The control animals were wild-type N2 strain. SAA-ablated animals were transgenic strain (P*lad-2*::Cre; P*lim-4*::loxP::PH-miniSOG). Ctrl, low viscosity: n = 233, 11 animals; Ctrl, middle viscosity: n = 239, 16 animals; Ctrl, high viscosity: n = 198, 11 animals; SAA-ablated, low viscosity: 190, 22 animals; SAA-ablated, middle viscosity: n = 224, 13 animals; SAA-ablated, high viscosity: n = 80, 16 animals. **E**. Representative curvature kymographs of control and SAA-ablated animals during forward movements. Body curvature was normalized by the worm body length *L* into a dimensionless unit *κ* · *L*. The worms were swimming in 25% dextran M9 solution.

**Fig. S3.**
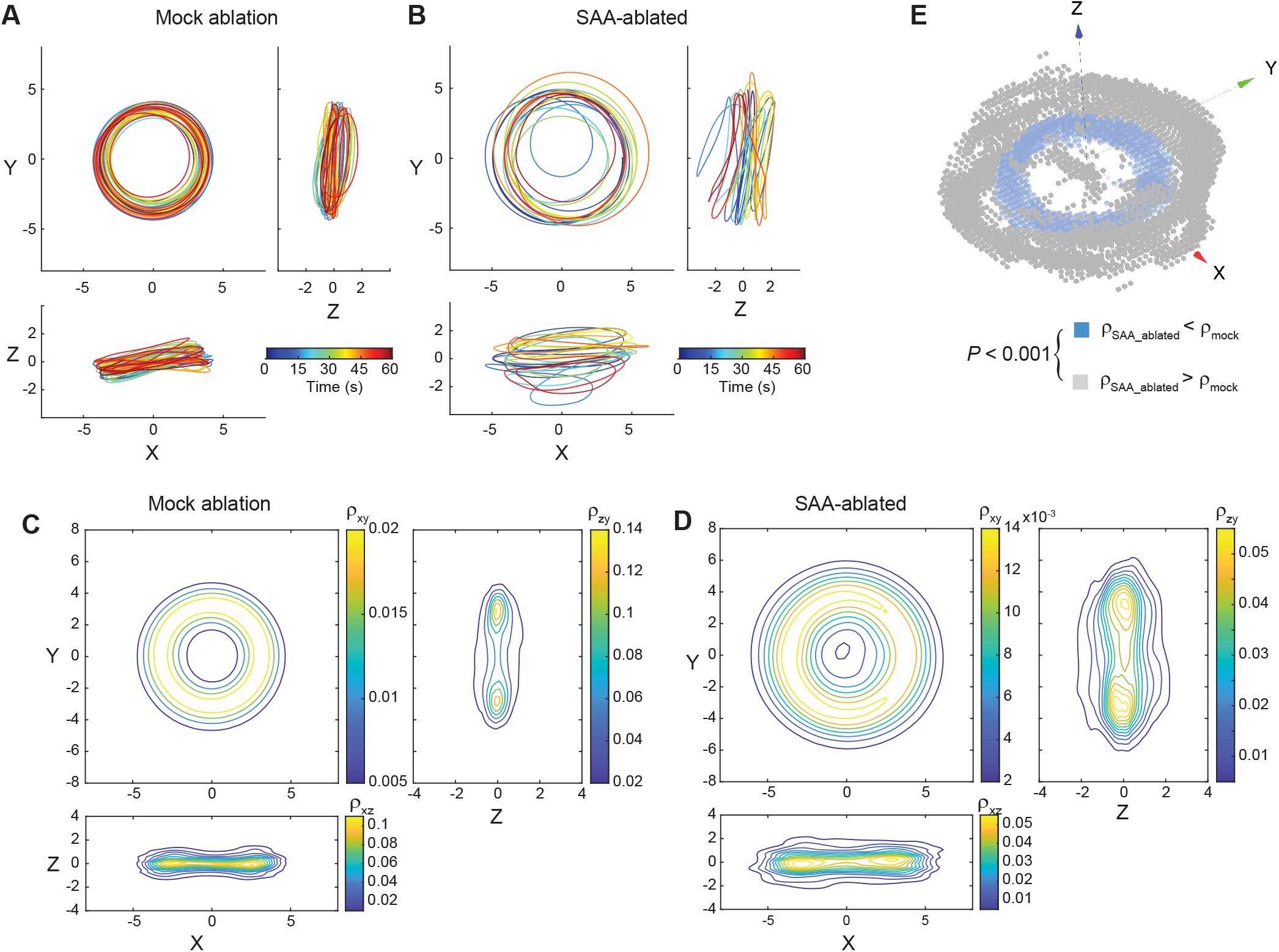
SAA stabilize rhythmic motion in forward movements of swimming worms. **A, B**. Time evolution of a phase trajectory in a control animal (**A**) and an SAA-ablated animal (**B**). Time is color-coded on the phase curves. **C**. Density of trajectories embedded in 3-dimensional phase space during forward movements of control animals. Each of the three sub-panels represents a density projection onto a plane spanned by two orthogonal directions. **D**. Similar to **C**, but for SAA-ablated animals. **E**. Local density differences between **C** and **D** were visualized by a voxelgram (*Phase space reconstruction and analysis*). The experiments in this figure were carried out in 25% dextran M9 solution, using strain P*lad-2*::Cre; P*lim-4*::loxP::PH-miniSOG. Ctrl: n=96, 12 animals; SAA-ablated: n =108, 9 animals.

**Fig. S4.**
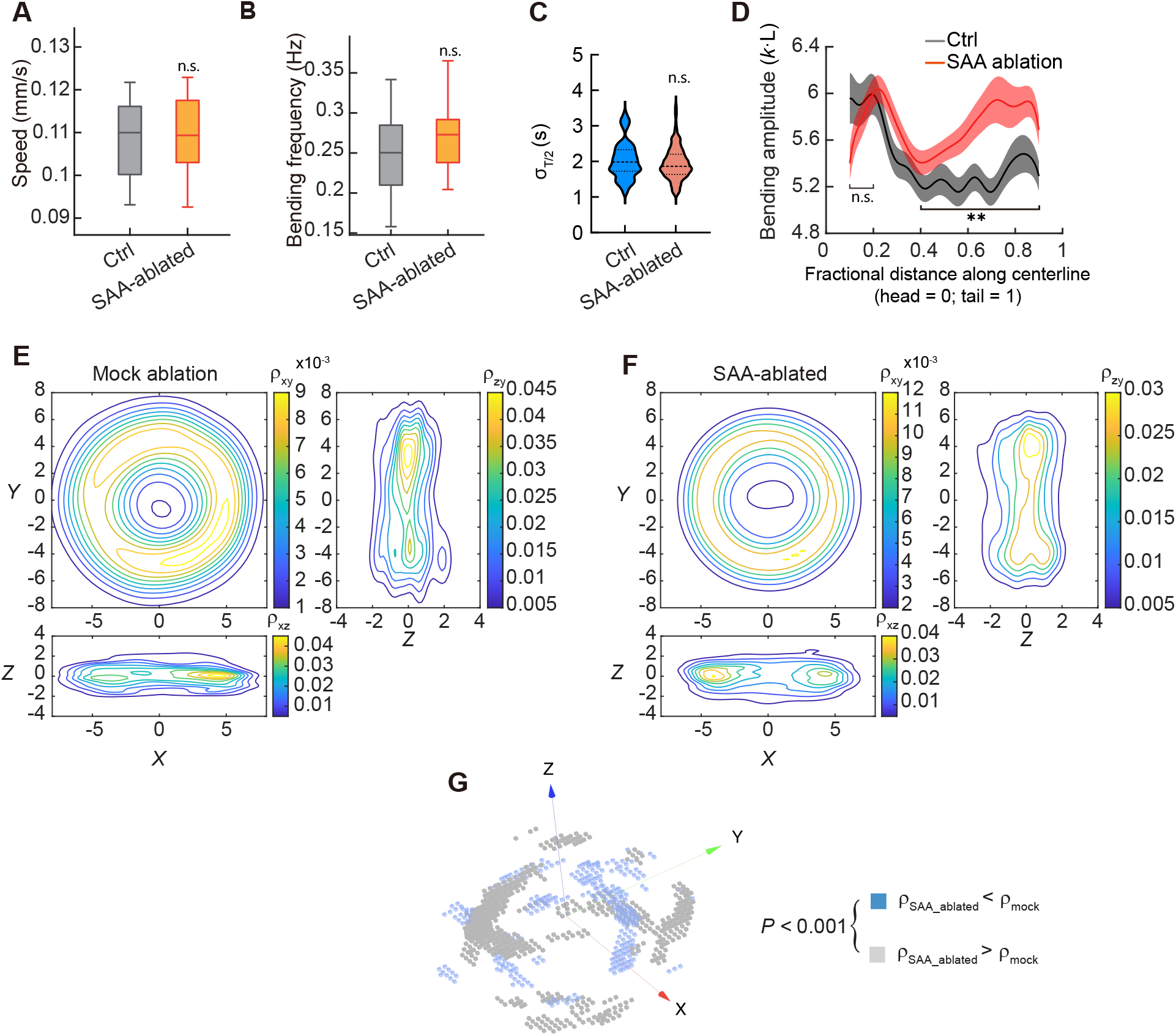
Kinematic parameters of SAA-intact and SAA-ablated worms with a similar speed distribution. **A-C**. We sub-sampled control animals (SAA mock ablated) and SAA-ablated animals with a similar speed distribution to compare the fast kinematic parameters in forward locomotion. We compared forward movement velocity, Undulation frequency and standard deviation of the half-cycle duration. n.s. p *>* 0.05, two-sample t-test with Welch’s correction. Box plots show the quartiles of the dataset, with whiskers extending from minimum to maximum, dashes show mean. **D**. Whole body bending amplitude for both control and SAA-ablated animals. The line indicates the average; the shaded region represents the SEM across trials. **P = 0.001, fractional distance ∈ [0.4, 0.9]; Mann–Whitney U test was performed on the spatially averaged bending amplitude across the body of the worm (Methods). **E**. Density of trajectories embedded in 3-dimensional phase space during forward movements of control animals. Each of the three sub-panels represents a density projection onto a plane spanned by two orthogonal directions. **F**. Similar as **E**, but for SAA-ablated animals. **G**. Local density differences between **E** and **F** were visualized by a voxelgram (*Phase space reconstruction and analysis*). Ctrl: n=96, 12 animals; SAA-ablated: n =108, 9 animals. Both the control group and the actual SAA-ablated group consist of transgenic animals (Plad-2::Cre; Plim-4::loxP::PH-miniSOG). Ctrl: n = 27, 5 animals; SAA-ablated: n = 28, 8 animals.

**Fig. S5.**
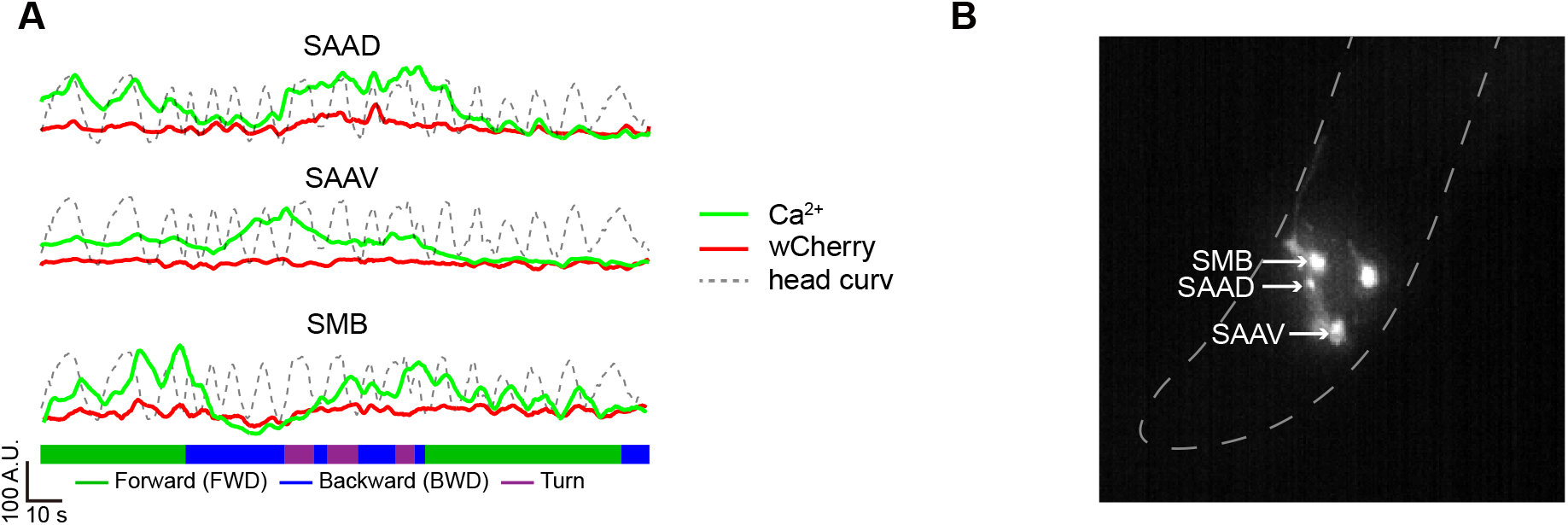
Ca^2+^ activity of SAA and SMB in a crawling worm. **A**. Raw fluorescence traces in the green Ca^2+^ indicator (wNEMOs) and the red reference channel (wcherry) observed while the worm was crawling on a 2% agarose pad covered with glass slide. Related to fig2 (**G**). **B**. Image of identified SAAD, SAAV, and SMB neurons under wide-field fluorescence microscopy.

**Fig. S6.**
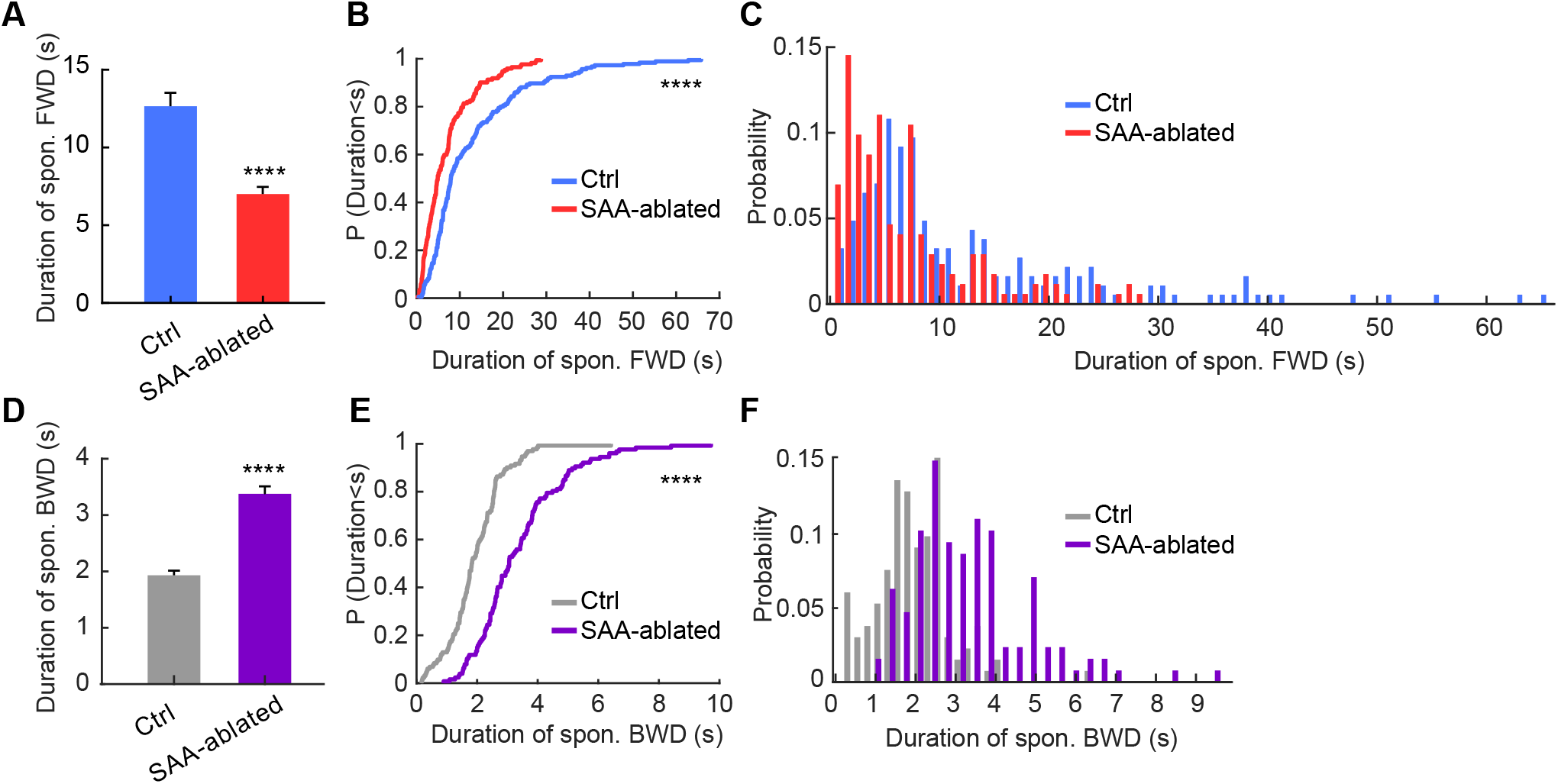
SAA contribute to organizing long timescale in swimming behaviors. **A**. Mean duration of forward movements. ****P *<* 0.0001, Mann–Whitney U test. Error bars represent SEM. Ctrl: n = 185, 11 animals; SAA-ablated: n = 172, 16 animals. **B**. Cumulative distributions of forward run length. Related to **A**. ****P *<* 0.0001, two-sample Kolmogorov–Smirnov test. **C**. Probability distributions of forward run length. Related to **A. D**. Mean duration of spontaneous reversals. ****P *<* 0.0001, Mann–Whitney U test. Error bars represent SEM. The control animals were N2 strain. SAA-ablated animals were transgenic strain (P*lad-2*::Cre; P*lim-4*::loxP::PH-miniSOG). Ctrl: n = 132, 11 animals; SAA-ablated: n = 127, 16 animals. **E**. Cumulative distributions of spontaneous reversal length. Related to **D**. ****P *<* 0.0001, two-sample Kolmogorov–Smirnov test. **F**. Probability distributions of reversal length. Related to **D**. All animals swam in a viscous solution (800 mPa·s).

**Fig. S7.**
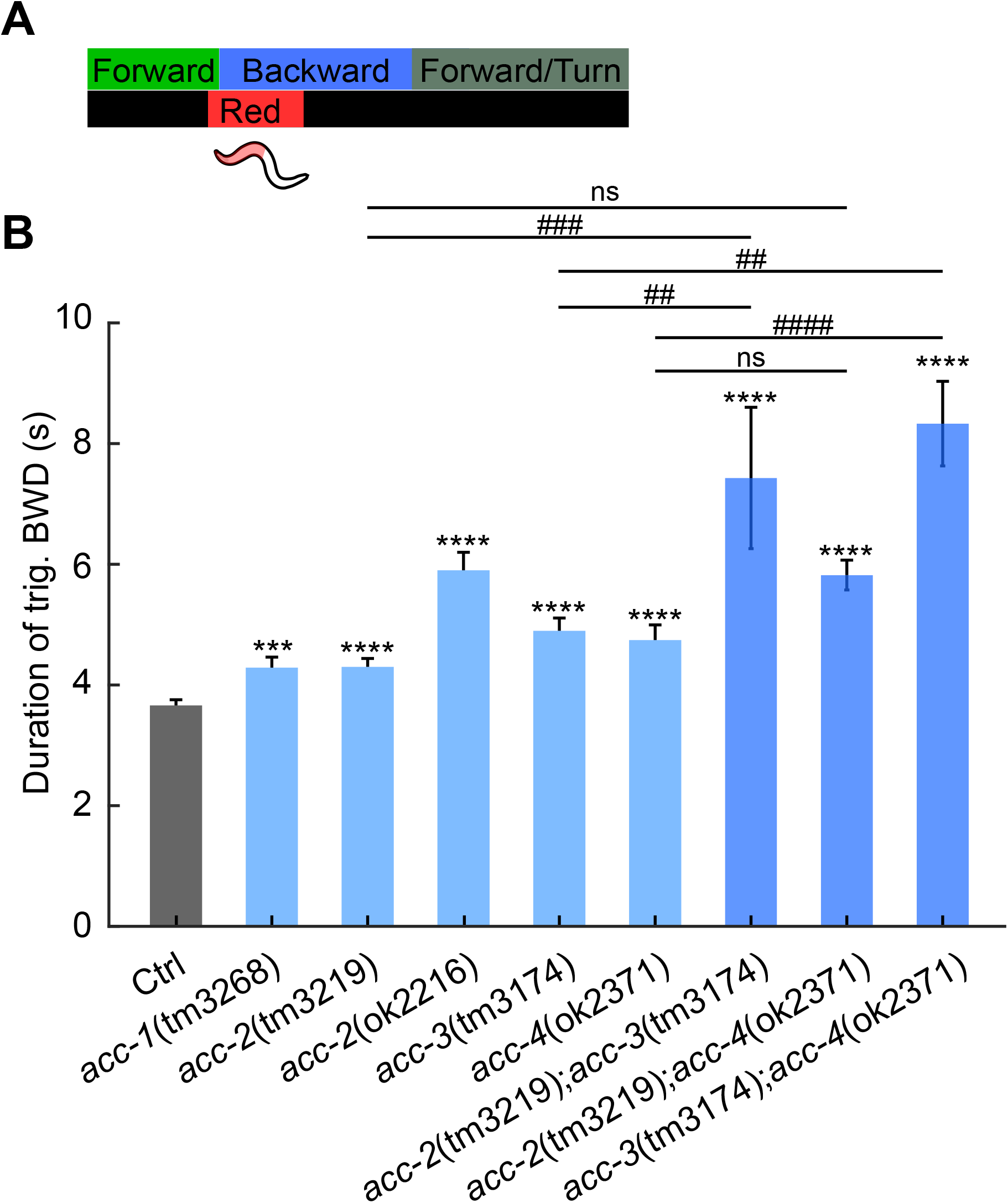
Acetylcholine-gated chloride channel subunits function synergistically to modulate reversal length during ALM/AVM-triggered escape responses. The duration of ALM/AVM-triggered reversals in a single *acc* mutant or a double mutant. ***P*<*0.001, ****P *<* 0.0001, compared with control; ##P*<*0.01, ###P*<*0.001, ####P*<*0.0001, compared between corresponding mutants; Mann–Whitney U test with with Bonferroni correction. Error bars represent SEM. Ctrl (N2): n = 414, 61 animals; *acc-1(tm3268)*: n = 114, 16 animals; *acc-2(tm3219)*: n = 100, 10 animals; *acc-2(ok2216)*: n = 88, 12 animals; *acc-3(tm3174)*: n = 97, 11 animals; *acc-4(ok2371)*: n = 123, 16 animals; *acc-2(tm3219)*; *acc-3(tm3174)*: n = 24, 4 animals; *acc-2(tm3219)*; *acc-4(ok2371)*: n = 134, 20 animals; *acc-3(tm3174)*; *acc-4(ok2371)*: n = 171, 30 animals.

**Fig. S8.**
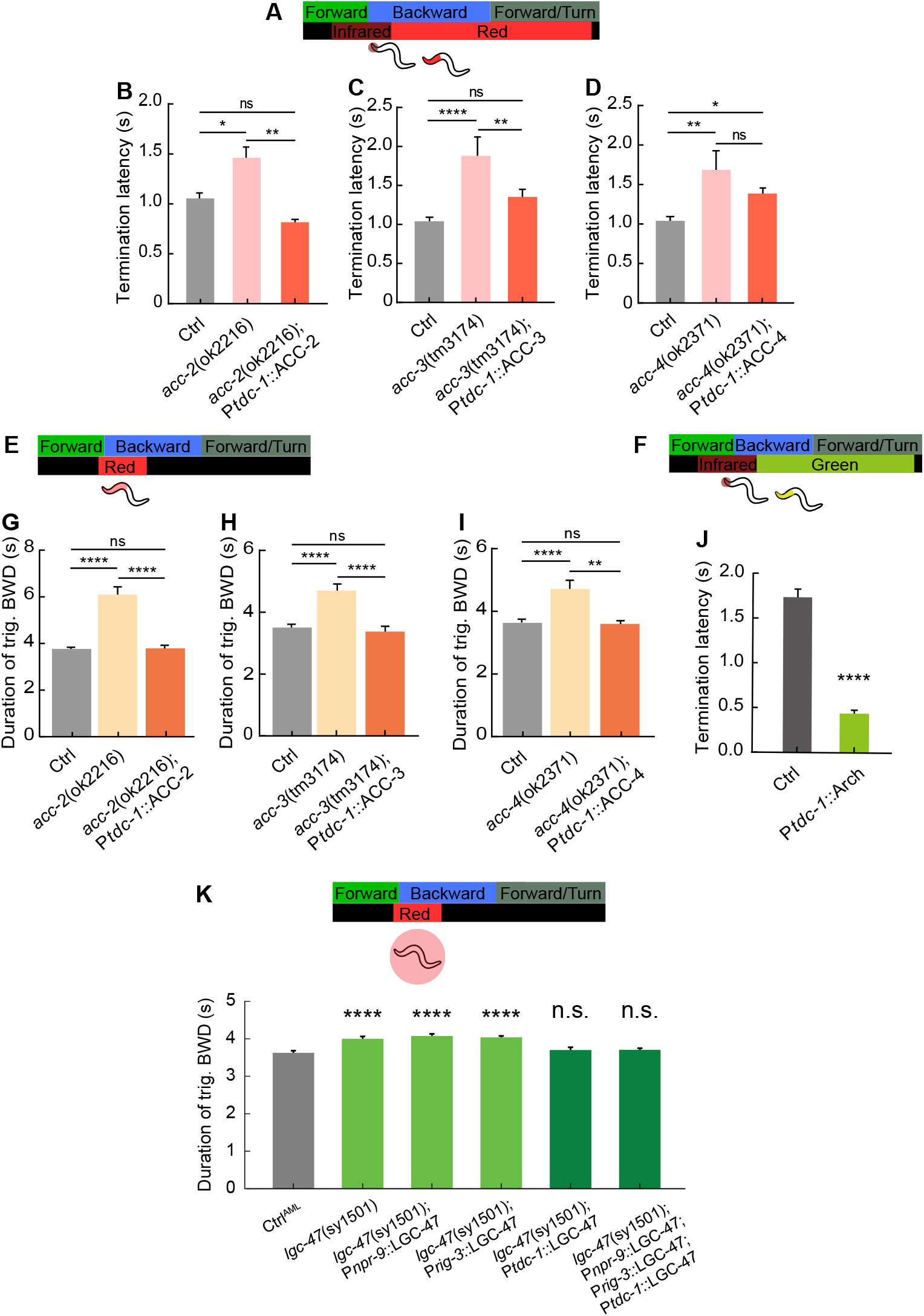
RIM communicates with SAA to terminate reversals via inhibitory acetylcholine synapses. **A**. Schematic experimental procedure for activation of SAA*/*RIV*/*SMB during thermally induced escape responses, same as fig. 5B. Related to **B, C**, and **D. B**. Termination latency in control, *acc-2* mutant, as well as animals in which ACC-2 was specifically restored in RIM. Ctrl: n = 229, 53 animals; *acc-2(ok2216)*: n = 103, 22 animals; *acc-2(ok2216)*;P*tdc-1::ACC-1*: n = 104, 19 animals. **C**. Termination latency in control, *acc-3* mutant, as well as animals in which ACC-3 was specifically restored in RIM. Ctrl: n = 229, 53 animals; *acc-3(tm3174)*: n = 126, 24 animals; *acc-3(tm3174)*;P*tdc-1::ACC-1*: n = 78, 14 animals. **D**. Termination latency in control, *acc-4* mutant, as well as animals in which ACC-4 was specifically restored in RIM. Ctrl: n = 229, 53 animals; *acc-4(ok2371)*: n = 141, 28 animals; *acc-4(ok2371)*;P*tdc-1::ACC-1*: n = 143, 26 animals. **E**. Schematic experimental procedure for triggering escape responses, same as fig. 4H. Optogenetic stimulation would activate AVM/ALM mechanosensory neurons. Related to **G, H**, and **I. G**. Duration of AVM/ALM-triggered reversal of control group, *acc-2* mutant and animals in which ACC-2 were specifically rescued in RIM. Ctrl: n = 414, 61 animals; *acc-2(ok2216)*: n = 88, 12 animals; *acc-2(ok2216)*;P*tdc-1::ACC-1*: n = 112, 19 animals. **H**. Duration of AVM/ALM-triggered reversal of control group, *acc-3* mutant and animals in which ACC-3 were specifically rescued in RIM. Ctrl: n = 414, 61 animals; *acc-3(tm3174)*: n = 88, 12 animals; *acc-3(tm3174)*;P*tdc-1::ACC-1*: n = 97, 11 animals. **I**. Duration of AVM/ALM-triggered reversal of control group, *acc-4* mutant and animals in which ACC-4 were specifically rescued in RIM. Ctrl: n = 414, 61 animals; *acc-4(ok2371)*: n = 123, 16 animals; *acc-4(ok2371)*;P*tdc-1::ACC-1*: n = 127, 21 animals. **F**. Schematic experimental procedure for inhibition of RIM during thermally induced escape responses. Reversal was triggered by an infrared laser (1480 nm, 50 mW/mm^2^) focusing on the worm head for 1 s, followed by 7 s green light optogenetic inhibition. Related to **J. J**. Termination latency between the onset of inhibiting RIM and the end of a reversal. Ctrl: n = 118, 19 animals; RIM::Arch: n = 85, 18 animals. **K**. Duration of all-TRN-triggered reversal in Ctrl, lgc-47 mutant, as well as animals in which LGC-47 was specifically restored in AIB, AVA, RIM, and all three neurons simultaneously. The number of optogenetic stimulus events from left to right are 550, 498, 622, 1280, 341, and 889. All statistical tests: *P *<* 0.05, **P *<* 0.01, ****P *<* 0.0001, Mann–Whitney U test or Mann-Whitney U test with Bonferroni correction. Error bars represent SEM.

**Fig. S9.**
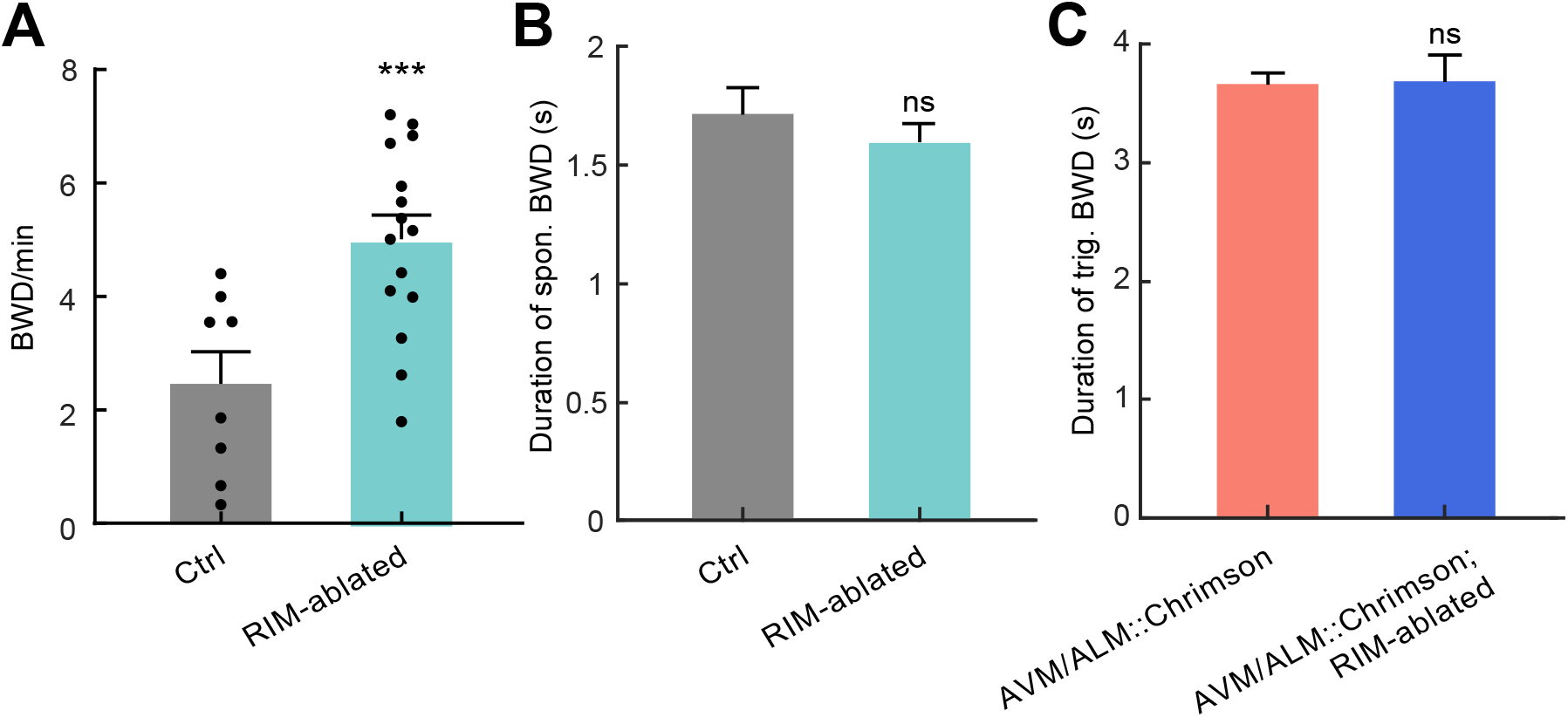
RIM-ablated animals make more frequent spontaneous reversals. **A**. Reversal frequency of control and RIM-ablated animals. Black dot represents an individual animal. Ctrl: n = 8 animals; RIM-ablated: n = 15 animals. **B**. Duration of spontaneous reversals in control and RIM-ablated animals are not significantly different. Ctrl: n = 112, 8 animals; RIM-ablated: n = 450, 15 animals. **C**. Duration of ALM/AVM triggered reversals in control and RIM-ablated animals are not significantly different. Ctrl: n = 414, 61 animals; RIM-ablated: n = 90, 17 animals. All statistical tests: Mann–Whitney U test, ***P *<* 0.001, error bars represent SEM.

**Fig. S10.**
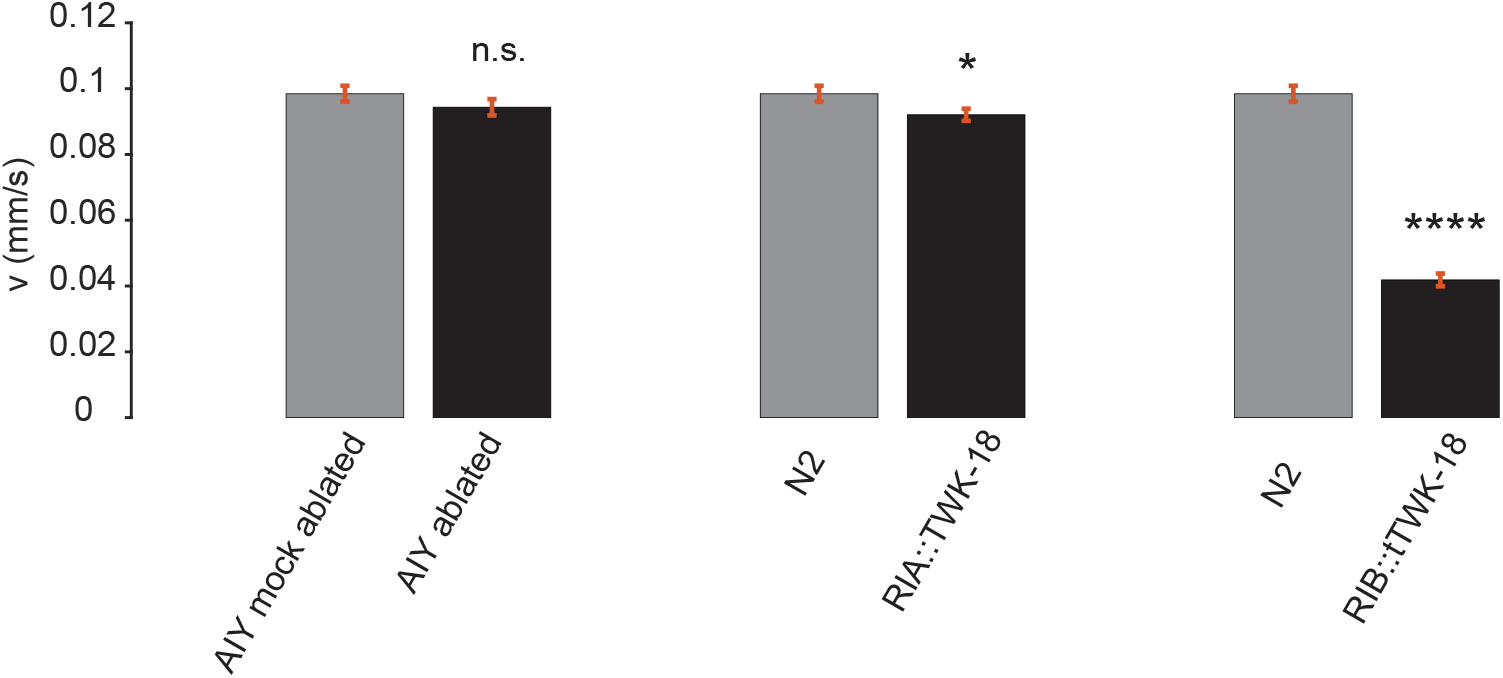
Influence of interneurons on forward movement speed. The error bars represents SEM. Bootstrap statistical test was performed on the mean speed between mocked-ablated and AIY-ablated (*Pttx-3::PH-miniSOG*) animals (n = 70, n = 72), N2 and RIA silenced animals (n = 70, n = 90), N2 and RIB silenced animals (n = 70, n = 46). n.s., P = 0.25, *P = 0.045, ****P *<* 0.0001. n is the number of tracks.

**Fig. S11.**
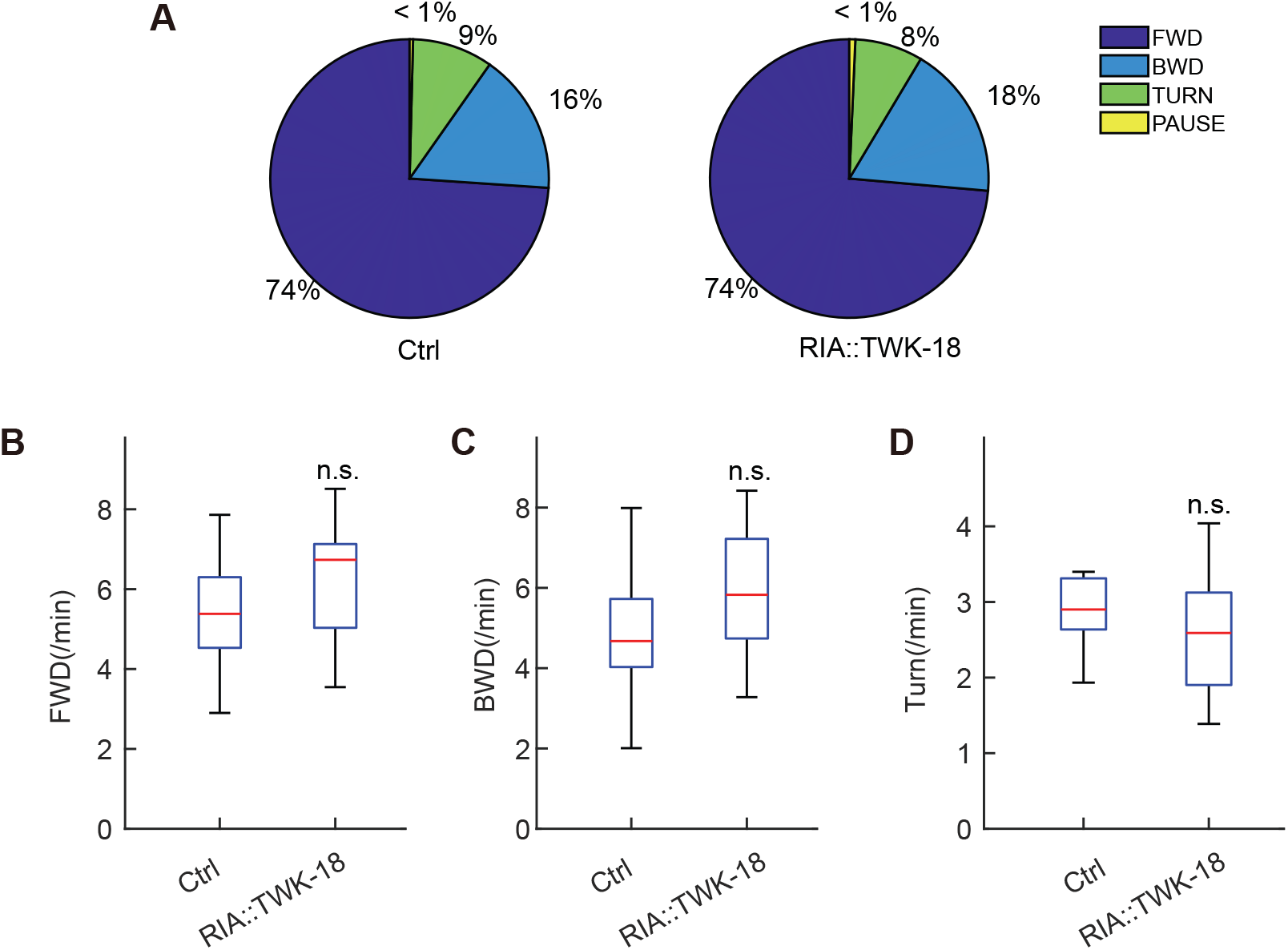
RIA does not contribute to organizing long timescale behaviors. **A**. The percentage of time spent on forward movements, reversals, turns, or pauses. The control group represents wild type animals. **B-D**. Mean occurrences per minute of spontaneous forward movements, reversals and turns in control and RIA::TWK-18 animals. Mann–Whitney U test. Ctrl: n = 11 animals; RIA::TWK-18(gf): n = 10 animals. Red lines represent the mean, and error bars represent the data range.

**Fig. S12.**
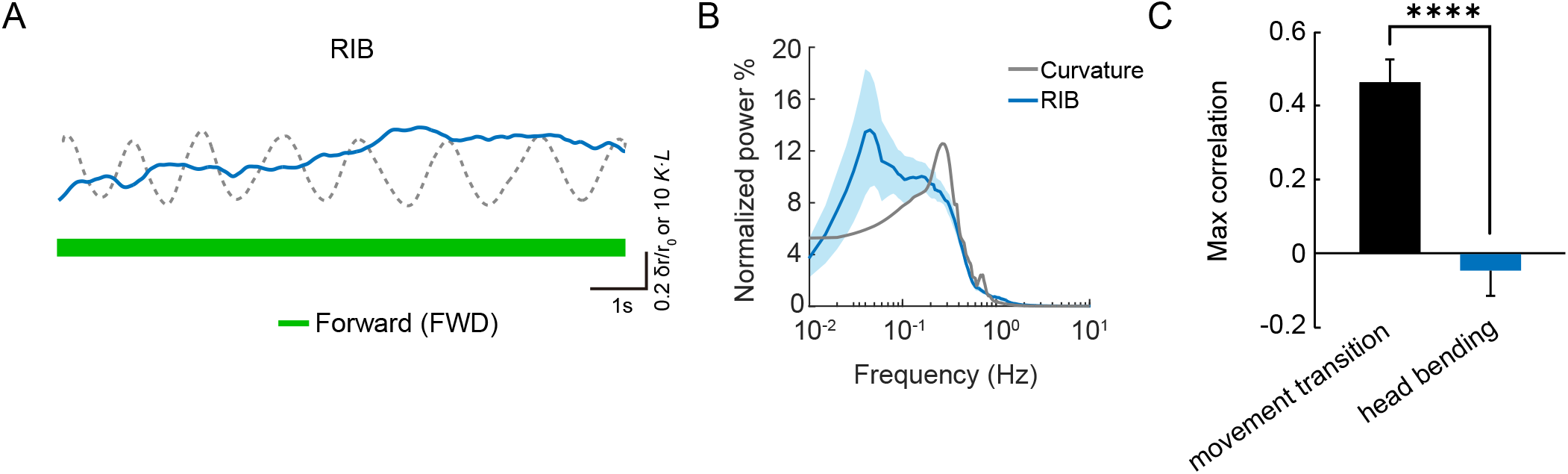
Calcium activity of RIB in crawling worms. **A**. A representative trace displays RIB activity alongside the curvature of the worm’s head during forward locomotion. A blue trace illustrates changes in the GCaMP6 to mCardinal ratio, while a dark dashed trace indicates the dynamics of head curvature ([0.1, 0.2] fractional distance along the worm). **B**. Average power spectral density of RIB during forward locomotion. The shaded regions represent SD. **C**. Maximum correlation between neuronal activity and head curvature during forward locomotion, as well as between neuronal activity and backward-to-forward state transition events. RIB activity showed a strong correlation with backward-to-forward state transition (also see reference 39 in the main text), but not with head bending curvature during forward locomotion. ****p *<* 0.0001, two-sample t-test. n = 30 trials from 5 worms.

#### Supplementary tables (S2) Information of plasmids and strains

**Table S1.** Plasmids used in this work.

| Identifier | Plasmid |
| --- | --- |
| quan0713 | *Ptdc-1::Chrimson::mCherry* |
| quan0084 | *Ptdc-1::Arch::GFP* |
| quan0624 | *Ptdc-1::wCherry* |
| quan0495 | *Ptdc-1::tomm-20::miniSOG::wCherry* |
| quan0198 | *Plim-4(-3328-2174)v1p1::Chrimson::mCherry* |
| quan0160 | *Plim-4(-3328-2174)v1p1::tomm-20::miniSOG::wCherry* |
| quan0197 | *Plim-4(-3328-2174)v1p1::GCaMP6::wCherry* |
| quan0451 | *Plim-4(-3328-2174)v1p1::TeTx::wCherry* |
| quan0047 | *Pmec-4::Chrimson::mCherry* |
| quan0657 | *Ptdc-1::ACC-2(gene)::SL2::GFP* |
| quan0659 | *Ptdc-1::ACC-4(cDNA)::SL2::GFP* |
| quan0660 | *Ptdc-1::ACC-1(gene)::SL2::GFP* |
| quan0711 | *Ptdc-1::ACC-3(cDNA)::SL2::GFP* |
| quan0598 | *Pacc-3::GFP* |
| quan0599 | *Pacc-1::GFP* |
| quan0603 | *Pacc-2::GFP* |
| quan0633 | *Pacc-4::GFP* |
| quan0602 | *Pnmr-1::wCherry* |
| quan0051 | *Plin-44::GFP* |
| quan0232 | *Punc-122::RFP* |
| quan0134 | *Plin-44::mCherry* |
| quan0421 | *Plad-2::Cre* |
| quan0384 | *Plim-4(-3328-2174)v1p1::loxP::PBSTOP::loxP::Arch::wCherry* |
| quan0729 | *Plim-4(-3328-2174)v1p1::loxP::PBSTOP::loxP::tomm-20::miniSOG::wCherry* |
| quan0385 | *Plim-4(-3328-2174)v1p1::loxP::PBSTOP::loxP::Chrimson::wCherry* |
| quan0199 | *Plim-4(-3328-2174)v1p1::Arch::wCherry* |
| quan0918 | *Plim-4(-3328-2174)v1p1::wNEMOs* |
| quan0040 | *Psto-3::GCaMP6::2\*NLS::mCardinal* |
| quan0488 | *Pglr-3::TWK-18(gf)::wCherry* |
| RRID: Addgene_107745 | *Pmec-4::Chrimson::SL2::mCherry* |
| RRID: Addgene_225923 | *Pnpr-9::Al::LGC-47::SL2::tagBFP* |
| RRID: Addgene_225924 | *Prig-3::Al::LGC-47::SL2::GFP* |
| RRID: Addgene_225925 | *Ptdc-1::Al::LGC-47::SL2::his-24::tagRFP* |

**Table S2.**
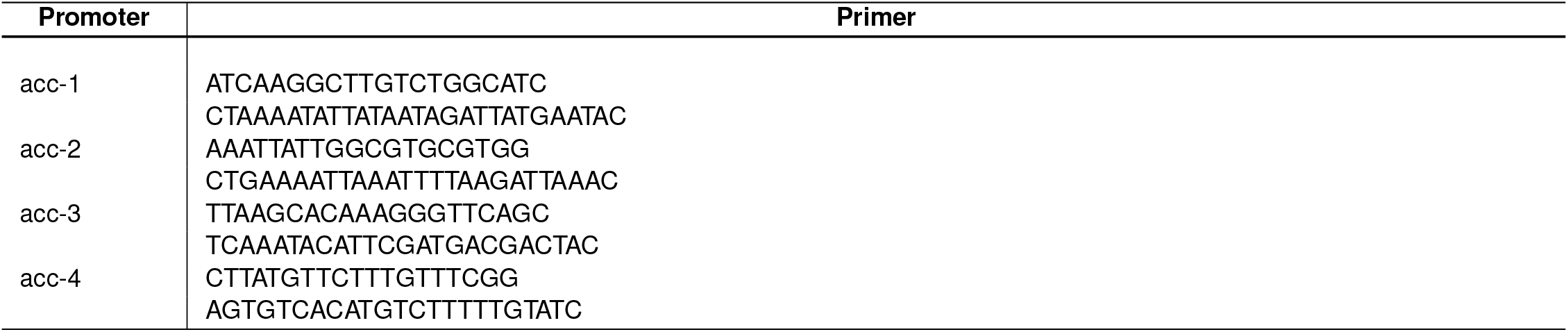
Primers used in this work.

**Table S3.** Strains used in this work.

| Identifier | Strain |
| --- | --- |
| WEN0902 | wenls1009[P<sub>lim-4</sub>(-3328-2174)v1p1::Chrimson::mCherry(30 ng/μL); P<sub>lin-44</sub>::GFP(10 ng/μL)];<br>wenEx0902[P<sub>tdc-1</sub>::PH-miniSOG::mCherry(30 ng/μL); P<sub>unc-122</sub>::RFP(30 ng/μL)] |
| WEN0920 | acc-1(tm3268)IV; wenls1015[P<sub>mec-4</sub>::Chrimson::mCherry; P<sub>lin-44</sub>::GFP]; wenEx0920[P<sub>tdc-1</sub>::ACC-1::GFP(20ng/μL);<br>P<sub>lin-44</sub>::mCherry(10 ng/μL)] |
| WEN0921 | acc-2(tm3219)IV; wenls1015[P<sub>mec-4</sub>::Chrimson::mCherry; P<sub>lin-44</sub>::GFP]; wenEx0921[P<sub>tdc-1</sub>::ACC-2::GFP(20 ng/μL);<br>P<sub>lin-44</sub>::mCherry(10 ng/μL)] |
| WEN0922 | acc-2(ok2216)IV; wenls1015[P<sub>mec-4</sub>::Chrimson::mCherry; P<sub>lin-44</sub>::GFP]; wenEx0922[P<sub>tdc-1</sub>::ACC-2::GFP(20 ng/μL);<br>P<sub>lin-44</sub>::mCherry(10 ng/μL)] |
| WEN0923 | acc-3(tm3174)X; wenls1015[P<sub>mec-4</sub>::Chrimson::mCherry; P<sub>lin-44</sub>::GFP]; wenEx0923[P<sub>tdc-1</sub>::ACC-3::GFP(20 ng/μL);<br>P<sub>lin-44</sub>::mCherry(10 ng/μL)] |
| WEN0924 | acc-4(ok2371)III; wenls1015[P<sub>mec-4</sub>::Chrimson::mCherry; P<sub>lin-44</sub>::GFP]; wenEx0924[P<sub>tdc-1</sub>::ACC-4::GFP(20 ng/μL);<br>P<sub>lin-44</sub>::mCherry(10 ng/μL)] |
| WEN0925 | acc-1(tm3268)IV; wenls1009[P<sub>lim-4</sub>(-3328-2174)v1p1::Chrimson::mCherry; P<sub>lin-44</sub>::GFP] |
| WEN0926 | acc-2(tm3219)IV; wenls1009[P<sub>lim-4</sub>(-3328-2174)v1p1::Chrimson::mCherry; P<sub>lin-44</sub>::GFP] |
| WEN0927 | acc-2(ok2216)IV; wenls1009[P<sub>lim-4</sub>(-3328-2174)v1p1::Chrimson::mCherry; P<sub>lin-44</sub>::GFP] |
| WEN0928 | acc-3(tm3174)X; wenls1009[P<sub>lim-4</sub>(-3328-2174)v1p1::Chrimson::mCherry; P<sub>lin-44</sub>::GFP] |
| WEN0929 | acc-4(ok2371)III; wenls1009[P<sub>lim-4</sub>(-3328-2174)v1p1::Chrimson::mCherry; P<sub>lin-44</sub>::GFP] |
| WEN1009 | wenls1009[P<sub>lim-4</sub>(-3328-2174)v1p1::Chrimson::mCherry; P<sub>lin-44</sub>::GFP]; 4* out-cross] |
| WEN1015 | wenls1015[P<sub>mec-4</sub>::Chrimson::mCherry; P<sub>lin-44</sub>::GFP]; 4* out-cross] |
| WEN1025 | acc-1(tm3268)IV; wenls1015[P<sub>mec-4</sub>::Chrimson::mCherry; P<sub>lin-44</sub>::GFP] |
| WEN1026 | acc-2(tm3219)IV; wenls1015[P<sub>mec-4</sub>::Chrimson::mCherry; P<sub>lin-44</sub>::GFP] |
| WEN1027 | acc-2(ok2216)IV; wenls1015[P<sub>mec-4</sub>::Chrimson::mCherry; P<sub>lin-44</sub>::GFP] |
| WEN1028 | acc-3(tm3174)X; wenls1015[P<sub>mec-4</sub>::Chrimson::mCherry; P<sub>lin-44</sub>::GFP] |
| WEN1030 | acc-4(ok2371)III; wenls1015[P<sub>mec-4</sub>::Chrimson::mCherry; P<sub>lin-44</sub>::GFP] |
| WEN1059 | acc-4(ok2371)III; acc-2(tm3219)IV; wenls1015[P<sub>mec-4</sub>::Chrimson::wCherry; P<sub>lin-44</sub>::GFP] |
| WEN1060 | acc-4(ok2371)III; acc-3(tm3174)X; wenls1015[P<sub>mec-4</sub>::Chrimson::wCherry; P<sub>lin-44</sub>::GFP] |
| WEN1062 | acc-2(tm3219)IV; acc-3(tm3174)X; wenls1015[P<sub>mec-4</sub>::Chrimson::wCherry; P<sub>lin-44</sub>::GFP] |
| WEN1064 | wenEx1064[P<sub>lim-4</sub>(-3328-2174)v1p1::PH-miniSOG::UrSL::wCherry(20 ng/μL); P<sub>unc-122</sub>::RFP(10 ng/μL)] |
| WEN1065 | wenls1009[P<sub>lim-4</sub>(-3328-2174)v1p1::Chrimson::mCherry; P<sub>lin-44</sub>::GFP];<br>wenEx1065[P<sub>lim-4</sub>(-3328-2174)v1p1::PH-miniSOG::UrSL::wCherry(20 ng/μL); P<sub>unc-122</sub>::RFP(10 ng/μL)] |
| WEN0932 | wenEx0932[P<sub>tdc-1</sub>::Arch::GFP(30 ng/μL); P<sub>lin-44</sub>::mCherry(10 ng/μL)] |
| WEN0938 | wenEx0938[P<sub>lad-2</sub>::Cre(30 ng/μL); P<sub>lim-4</sub>(-3328-2174)v1p1::loxP::PBSTOP::loxP::Chrimson::wCherry (50 ng/μL);<br>P<sub>lin-44</sub>::mCherry(15 ng/μL)] |
| WEN0941 | acc-1(tm3268)IV; wenls1009[P<sub>lim-4</sub>(-3328-2174)v1p1::Chrimson::mCherry; P<sub>lin-44</sub>::GFP];<br>wenEx0941[P<sub>tdc-1</sub>::ACC-1::GFP(20 ng/μL); P<sub>lin-44</sub>::mCherry(10 ng/μL)] |
| WEN0942 | acc-2(ok2216)IV; wenls1009[P<sub>lim-4</sub>(-3328-2174)v1p1::Chrimson::mCherry; P<sub>lin-44</sub>::GFP];<br>wenEx0942[P<sub>tdc-1</sub>::ACC-2::GFP(20 ng/μL); P<sub>lin-44</sub>::mCherry(10 ng/μL)] |
| WEN0943 | acc-3(tm3174)X; wenls1009[P<sub>lim-4</sub>(-3328-2174)v1p1::Chrimson::mCherry; P<sub>lin-44</sub>::GFP];<br>wenEx0943[P<sub>tdc-1</sub>::ACC-3::GFP(20 ng/μL); P<sub>lin-44</sub>::mCherry(10 ng/μL)] |
| WEN0944 | acc-4(ok2371)III; wenls1009[P<sub>lim-4</sub>(-3328-2174)v1p1::Chrimson::mCherry; P<sub>lin-44</sub>::GFP];<br>wenEx0944[P<sub>tdc-1</sub>::ACC-4::GFP(20 ng/μL); P<sub>lin-44</sub>::mCherry(10 ng/μL)] |
| WEN0945 | acc-2(tm3219)IV; wenls1009[P<sub>lim-4</sub>(-3328-2174)v1p1::Chrimson::mCherry; P<sub>lin-44</sub>::GFP];<br>wenEx0945[P<sub>tdc-1</sub>::ACC-2::GFP(20 ng/μL); P<sub>lin-44</sub>::mCherry(10 ng/μL)] |
| WEN0950 | wenEx0950[P<sub>lad-2</sub>::Cre(30 ng/μL); P<sub>lim-4</sub>(-3328-2174)v1p1::loxP::PBSTOP::loxP::tomm-20::miniSOG::wCherry(50 ng/μL);<br>P<sub>unc-122</sub>::RFP(10 ng/μL)] |
| WEN0567 | wenEx0567[P<sub>npr-9</sub>::Chrimson::mCherry; P<sub>lin-44</sub>::GFP(10 ng/μL); P<sub>lim-4</sub>(-3328-2174)v1p1::GCaMP6::wCherry] |
| WEN1053 | wenEx1053[P<sub>nmr-1</sub>::mCherry(30 ng/μL); P<sub>acc-1</sub>::GFP(30 ng/μL)] |
| WEN1054 | wenEx1054[P<sub>nmr-1</sub>::mCherry(30 ng/μL); P<sub>acc-2</sub>::GFP(30 ng/μL)] |
| WEN1055 | wenEx1055[P<sub>tdc-1</sub>::mCherry(30 ng/μL); P<sub>acc-3</sub>::GFP(30 ng/μL)] |
| WEN1056 | wenEx1056[P<sub>tdc-1</sub>::mCherry(30 ng/μL); P<sub>acc-4</sub>::GFP(30 ng/μL)] |
| WEN0575 | wenEx0475[P<sub>lim-4</sub>(-3328-2174)v1p1::Chrimson::mCherry; P<sub>lin-44</sub>::GFP(10 ng/μL)];<br>wenEx0575[P<sub>lim-4</sub>(-3328-2174)v1p1::TeTx::UrSL::GFP(30 ng/μL); P<sub>unc-122</sub>::RFP(10 ng/μL)] |
| WEN0951 | wenEx0951[P<sub>lad-2</sub>::Cre(30 ng/μL); P<sub>lim-4</sub>(-3328-2174)v1p1::loxP::PBSTOP::loxP::tomm-20::miniSOG::wCherry(50 ng/μL);<br>P<sub>lin-44</sub>::mCherry(25 ng/μL)] |
| WEN1070 | wenEx1070[P<sub>lad-2</sub>::Cre(30 ng/μL); P<sub>lim-4</sub>(-3328-2174)v1p1::loxP::PBSTOP::loxP::Arch::wCherry(100 ng/μL);<br>P<sub>unc-122</sub>::GFP(10 ng/μL)] |
| WEN1196 | wenEx1196[P<sub>lim-4</sub>(-3328-2174)v1p1: wNEMOs(30 ng/μL); P<sub>lim-4</sub>(-3328-2174)v1p1:Arch::wCherry(30 ng/μL);<br>P<sub>lin-44</sub>::mCherry(10 ng/μL)] |
| WEN1174 | *acc-1(tm3268)IV*; *wenEx1074[P<sub>lad-2</sub>::Cre(30 ng/μL)*,<br>*Plim-4(-3328–2174)v1p1::loxp::PBSSTOP::loxp::Chrimson::wCherry*, *Plin-44::mCherry(10 ng/μL)*] |
| WEN1092 | *acc-1(tm3268)IV*; *wenEx1074[P<sub>lad-2</sub>::Cre(30 ng/μL)*, *Plim-4(-3328–2174)v1p1::loxp::PBSSTOP::loxp::Chrimson::wCherry*;<br>*Ptdc-1::ACC-1::GFP(15ng/ul)*; *Plin-44::mCherry(10 ng/μL)*] |
| WEN1179 | *wenEx1179[wenEx1179[P<sub>lad-2</sub>::Cre(30 ng/μL)*; *Plim-4(-3328–2174)v1p1::loxp::PBSTOP::loxp::Chrimson::wCherry(50*<br>*ng/μL)*; *Ptdc-1::tomm-20::miniSOG::mCherry(30 ng/μL)*] |
| WEN0216 | *wenEx0216[P<sub>glr-3</sub>::TWK-18(gf)::wCherry(30 ng/μL)*; *Plin-44::GFP(10 ng/μL)*] |
| WEN0172 | *wenIs0172[P<sub>sto-3</sub>::GCaMP6::2\*NLS::mCardinal(50 ng/μL)*; *Plin-44::GFP(10 ng/μL)*] |
| AML67 | *wtfls46[P<sub>mec-4</sub>::Chrimson::SL2::mCherry(40 ng/μL)*] |
| AML597 | *lgc-47(sy1501)X*; *wtfls46[P<sub>mec-4</sub>::Chrimson::SL2::mCherry(40 ng/μL)*] |
| AML614 | *lgc-47(sy1501)X*; *wtfls46[P<sub>mec-4</sub>::Chrimson::SL2::mCherry(40 ng/μL)*]; *wtfEX535*<br>*[Ptdc-1::Al::LGC-47::SL2::his-24::tagRFP(30ng/μL)*; *Coel::GFP (70 ng/μL)*] |
| AML617 | *lgc-47(sy1501)X*; *wtfls46[P<sub>mec-4</sub>::Chrimson::SL2::mCherry(40 ng/μL)*]; *wtfEX538*<br>*[Pnpr-9::Al::LGC-47::SL2::tagBFP(30ng/μL)*; *Coel::GFP(70 ng/μL)*] |
| AML618 | *lgc-47(sy1501)X*; *wtfls46[P<sub>mec-4</sub>::Chrimson::SL2::mCherry(40 ng/μL)*]; *wtfEX539 [Prig-3::Al::LGC-47::SL2::GFP(30*<br>*ng/μL)*; *Coel::GFP(70 ng/μL)*] |
| AM622 | *lgc-47(sy1501)X*; *wtfls46[P<sub>mec-4</sub>::Chrimson::SL2::mCherry(40 ng/μL)*]; *wtfEX543*<br>*[Ptdc-1::Al::LGC-47::SL2::his-24::tagRFP(30 ng/μL)*; *Pnpr-9::Al::LGC-47::SL2::tagBFP(30 ng/μL)*;<br>*Prig-3::Al::LGC-47::SL2::GFP(30 ng/μL)*; *Coel::GFP(70 ng/μL)*] |
| AM627 | *acc-1(tm3268)IV*; *wtfls46[P<sub>mec-4</sub>::Chrimson::SL2::mCherry(40 ng/μL)*] |

